# Targeted-Antibacterial-Plasmids (TAPs) combining conjugation and CRISPR/Cas systems achieve strain-specific antibacterial activity

**DOI:** 10.1101/2020.10.12.335968

**Authors:** Audrey Reuter, Cécile Hilpert, Annick Dedieu-Berne, Sophie Lematre, Erwan Gueguen, Guillaume Launay, Sarah Bigot, Christian Lesterlin

## Abstract

The global emergence of drug-resistant bacteria leads to the loss of efficacy of our antibiotics arsenal and severely limits the success of currently available treatments. Here, we developed an innovative strategy based on Targeted-Antibacterial-Plasmids (TAPs) that use bacterial conjugation to deliver CRISPR/Cas systems exerting a strain-specific antibacterial activity. TAPs are highly versatile as they can be directed against any specific genomic or plasmid DNA using the custom algorithm (CSTB) that identifies appropriate targeting spacer sequences. We demonstrate the ability of TAPs to induce strain-selective killing by introducing lethal double strand breaks (DSBs) into the targeted genomes. TAPs directed against a plasmid-born carbapenem resistance gene efficiently resensitise the strain to the drug. This work represents an essential step towards the development of an alternative to antibiotic treatments, which could be used for *in situ* microbiota modification to eradicate targeted resistant and/or pathogenic bacteria without affecting other non-targeted bacterial species.

## Introduction

The worldwide proliferation of drug-resistant bacteria is predicted to cause a dramatic increase in human deaths due to therapeutic failures in the next decades^1^. The constant emergence of bacterial resistances and the current low rate of antibiotic discovery emphasize the need to develop innovative antibacterial strategies that represent a real alternative to the use of antibiotics. Moreover, antibiotics generally lack specificity as they target processes that are essential to bacterial proliferation. Antibiotics consequently affect the whole treated bacterial community without discriminating between harmful and commensal strains, and lead to the population enrichment in drug-resistant strains.

Recent reports have demonstrated the possibility to achieve specific antimicrobial activity through the use of Clustered Regularly Interspaced Short Palindromic Repeats (CRISPR) and the associated Cas proteins. CRISPR/Cas systems can achieve bacterial killing through the induction of double-strand breaks (DSBs) to the chromosome by the Cas9 nuclease^2,3^. The expression of specific genes can also be inhibited through CRISPR interference (CRISPRi), when using the dead catalytic Cas9 enzyme (dCas9)^4,5^. CRISPR targeting relies on the ~16-20 nucleotide (nt) target-specific guide RNA (gRNA) sequence, which allow the recruitment of the Cas nuclease to the complementary DNA sequence^2,6^. Yet, to be used as practical antibacterial tools, CRISPR/Cas genes need to be delivered to the targeted bacterium. Bacterial DNA conjugation precisely offers the possibility to transfer long DNA segments to a range of bacterial species, with the transfer specificity depending on the considered conjugation system.

In this work, we present an innovative antibacterial methodology based on transferrable plasmids that carry CRISPR/Cas systems designed to induce antibacterial activity into specifically targeted recipient strains. These so-called Targeted-Antibacterial-Plasmids (TAPs) use the conjugation machinery (*tra* genes) encoded by a F plasmid contained in the donor strain to be transferred to *E. coli* strains and to closely related Gram-negative Enterobacteriaceae. The guide RNA (gRNA) sequence carried by the TAP determines the targeting of the antibacterial activity towards specific recipient strains only. Indeed, the CRISPR/Cas system will only be active in recipients that contain the DNA sequence complementary to the chosen gRNA sequence, while being inactive in other strains. TAPs can then easily be redirected against any bacterial species of interest by changing the gRNA sequence in one-step cloning. To identify strain-specific gRNA, we have developed a bio-informatic program CSTB (CRISPR Search Tool for Bacteria) that allows the rapid and robust identification of ~16-20 nt sequences on the basis of their presence or absence in the genome of bacterial strains selected on a phylogenetic tree. Here we demonstrate TAPs ability to induce efficient and strain-specific antibacterial activity *in vitro*.

## Results

### Targeted-Antibacterial-Plasmids (TAPs) modular design

TAPs derivate from the synthetic pSEVA plasmid collection^7^, and carry the pBBR1 origin of replication, a choice of resistant gene cassettes, and the *oriTF* origin of transfer of the F plasmid (Fig.1a). TAPs are consequently mobilizable by the conjugation machinery produced in *trans* from the conjugative F-Tn*10* helper plasmid contained in the donor cells (Fig.1b and Supplementary Fig.1a)^8,9^. We inserted the *Streptococcus pyogenes* wild-type *cas9* (for CRISPR activity) or catalytically dead *dcas9* gene (for CRISPRi activity) and the guide gRNA sequence composed of the constant tracrRNA scaffold and the target-specific crRNA spacer sequence (Fig.1a). Changing the crRNA spacer sequence in one-step-cloning allows reprogramming the targeting of the TAPs against any specific chromosome or plasmid DNA. Optionally, TAPs also carry either the *superfolder green fluorescent* (*sfgfp*) or the *mcherry* gene highly expressed from the broad-host range synthetic BioFab promoter^10^ to serve as plasmid transfer fluorescent reporter in microscopy and flow cytometry assays (Fig.1a).

**Figure 1.**
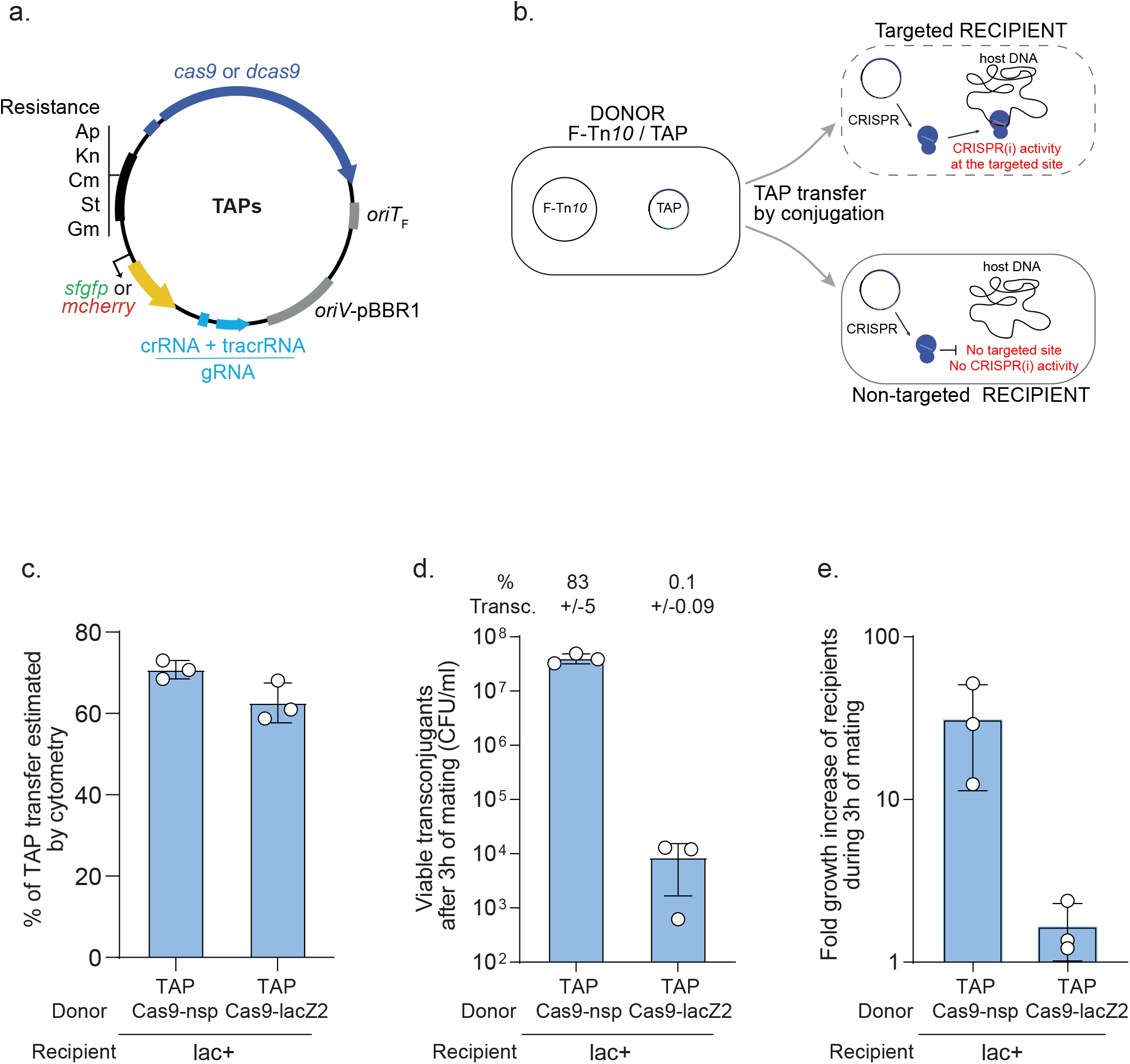
Transfer of TAP by F plasmid machinery mediates killing of a targeted *E. coli* strain. (**a**) TAPs modules consist of CRISPR system composed of wild type (*cas9*) or catalytically dead *cas9* (*dcas9*) genes expressed from the weak constitutive *BBa_J23107* promoter and a gRNA module expressed from the strong constitutive *BBa_J23119* promoter; the F plasmid origin of transfer (*oriT*F); the pBBR1 origin of replication (*oriV*), and a set of resistance cassettes (Ap, ampicillin; Kn, kanamycin; Cm, chloramphenicol; St, streptomycin; Gm, gentamycin), an optional cassette acrying the *sfgfp* or *mcherry* genes highly expressed from the broad-host range synthetic BioFab promoter (**b**) Diagram of the TAP antibacterial strategy. A donor strain produces the F plasmid conjugation machinery to transfer the TAP into the recipient strain. Targeted recipient carries sequence recognized by CRISPR(i) system that induces killing or gene expression inhibition. Non-targeted recipient lacking the spacer recognition sequence are insensitive to CRISPR(i) activity. (**c**) Histogram of TAPs transfer estimated by flow cytometry show that TAP_kn_-Cas9-nsp-GFP and TAP_kn_-Cas9-lacZ2-GFP are transferred with similar efficiency in recipient cells after 3h of mating. Donors TAP-Cas9-nsp (LY1371) or TAP-Cas9-lacZ2 (LY1380), recipient HU-mCherry lac+ (LY248). (**d**) Histograms of the concentration of viable transconjugants estimated by plating assays show viability loss associated with the acquisition of TAP-Cas9-lacZ2. The corresponding percentage of viable transconjugants (ratio T/R+T) is shown above each bar. (**e**) Fold-increase of the recipient population counts over the 3h of mating. Donors TAP-Cas9-nsp (LY1369) or TAP-Cas9-lacZ2 (LY1370), recipient lac+ (LY827). (**c-e**) Mean and SD are calculated from 3 independent experiments.

### Validation of TAPs CRISPR and CRISPRi activities

We addressed the ability of TAPs to induce efficient and specific Cas9-mediated killing (CRISPR) or dCas9-mediated gene expression inhibition (CRISPRi). First, TAPs ability to induce Cas9-mediated killing was confirmed using the previously described lacZ2 spacer that targets the *lacZ* gene of *E. coli*^11^. Transformation of the TAP-Cas9-lacZ2 plasmid into the lac+ MG1655 wt strain was ~1000-fold less efficient than in the isogenic lac- strain carrying a deletion of the targeted *lacZ* locus (Supplementary Fig.1b). By contract, the TAP-Cas9-nsp plasmid, which contains a non-specific (nsp) crRNA spacer that does not target *E. coli* genome, was transformed with equal efficiency in both lac+ and lac- strains (Supplementary Fig.1b). Second, TAPs ability to induce dCas9-mediated CRISPRi activity was validated by using the csgB spacer that targets the *csgB* promoter driving the production of cell-surface curli fimbriae^12^ in the MG1655 *E. coli* mutant strain OmpR234^13^. Congo Red (CR) staining on agar-plates and aggregation clumps formation in liquid medium were used as direct readouts for curli production^13,14^. The TAP-dCas9-csgB efficiently inhibits curli production by the OmpR234 strain, as reflected by the formation of white colony in the presence of CR and the inability to form aggregation clumps (Supplementary Fig.1c). By contrast, the non-specific TAP-dCas9-nsp add no effect on curli formation or aggregation in the OmpR234 strain. Besides, we confirmed that the constitutive production of the Cas9 or dCas9 from the TAPs do not cause growth defects (Supplementary Fig.1d) or elongated cell morphology (Supplementary Fig.1e), contrasting with the toxic effects reported in some systems^15–18^. These results demonstrate that TAPs ability to induce Cas9-mediated killing or dCas9-mediated gene expression inhibition is efficient and depends on the accurate targeting by the spacer sequence.

### TAPs-mediated killing of targeted recipient cells

Next, we addressed the ability of the TAPs to be transferred by conjugation and induce antibacterial activity in *E. coli* recipient cells. Conjugation was performed using the *E. coli* MG1655 donor strain that contains the F-Tn*10* helper plasmid and either the TAP-Cas9-nsp or TAP-Cas9-lacZ2 mobilizable plasmids. Using flow cytometry analysis, we quantified the transfer efficiency of these TAPs (carrying the sfGFP green fluorescent reporter) into a lac+ recipient strain that produces the red fluorescent histone-like protein HU-mCherry, encoded on the chromosome (Supplementary Fig.2a). Quantification of the transconjugants exhibiting combined red and green fluorescence show that TAP-Cas9-nsp and TAP-Cas9-lacZ2 are both transferred to ~65% of the recipient cell population after 3h of mating (Fig.1c and Supplementary Fig.2b). As expected, TAPs transfer requires the presence of the F-Tn*10* plasmid in the donor strain (Supplementary Fig.2c-e). Most importantly, the parallel plating of the conjugation mixes revealed a ~3.5-log decrease in the viability of TAP-Cas9-lacZ2 transconjugants compared to TAP-Cas9-nsp transconjugants (Fig.1d). This killing activity is also reflected by the lack of increase in the total recipient cells count during the three hours of mating with the TAP-Cas9-lacZ2 donor strain, compared to a ~20-fold increase with TAP-Cas9-nsp donors (Fig. 2e). Importantly, no killing effect is observed for either TAPs when using the isogenic lac- recipient strain lacking the targeted *lacZ* locus (Supplementary Fig.2f). These results show that TAP-Cas9-nsp and TAP-Cas9-lacZ2 are transferred with equal efficiency through the F-Tn*10* conjugation machinery. Yet, the acquisition of TAP-Cas9-lacZ2, but not TAP-Cas9-nsp, is associated with a loss of viability of the transconjugant cells.

**Figure 2.**
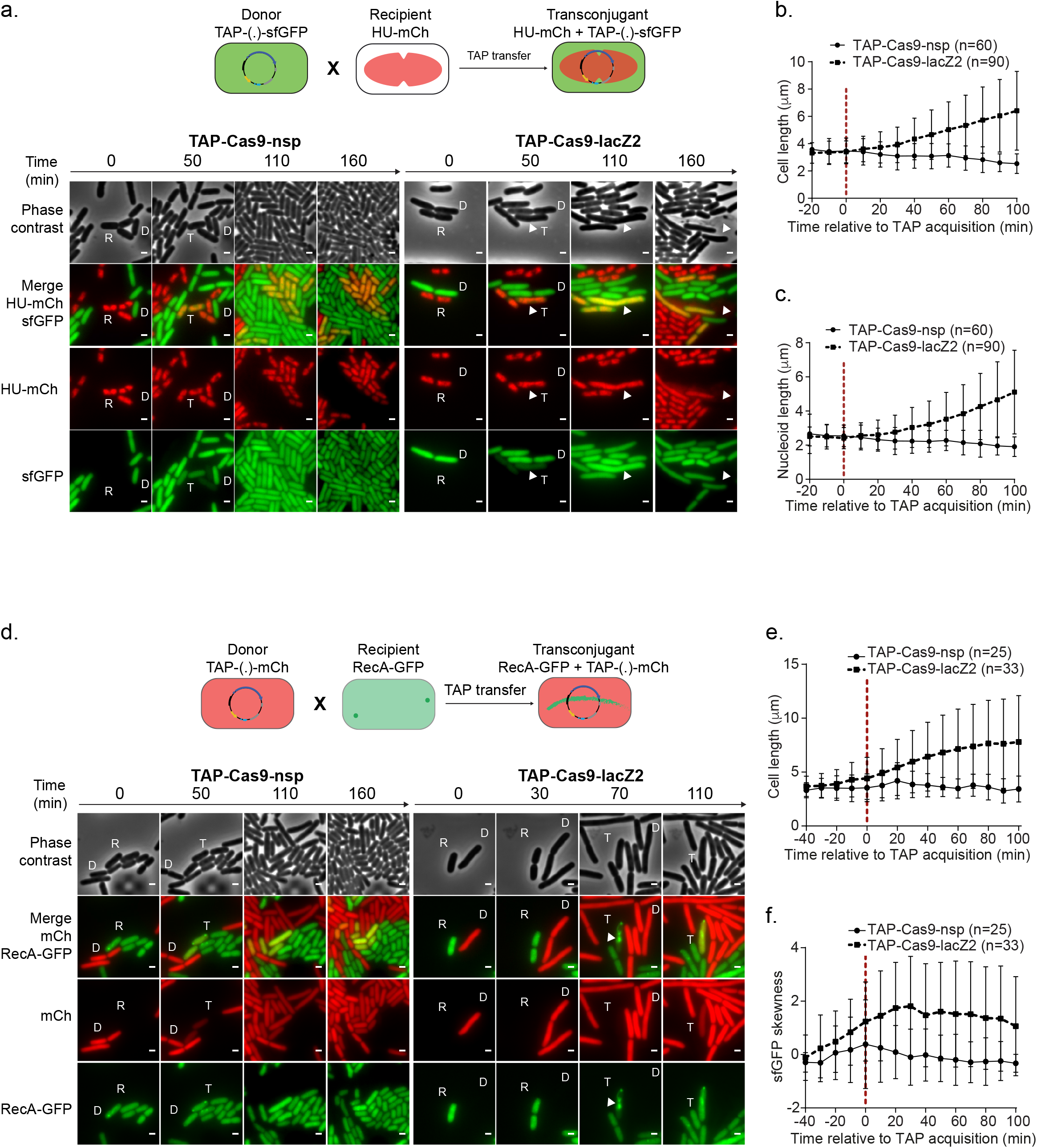
Real-time visualization of *E. coli* killing after acquisition of TAP. (**a**) Upper panel: diagram of the fluorescent reporter system allowing microcopy visualization of TAP transfer and subsequent nucleoid disorganization in transconjugants. Donor cells exhibit diffuse green fluorescence due to sfGFP production from TAP; HU-mCherry recipients exhibit red nucleoid associated fluorescence; transconjugants are identified by the production of both green and red fluorescence. Lower panel: time-lapse microscopy images performed in a microfluidic chamber. D (donor), recipient (R), and transconjugant (T) cells are indicated. Scale bar 1 μm. Donors TAP-Cas9-nsp (LY1371) or TAP-Cas9-lacZ2 (LY1380); recipient HU-mCherry lac+ (LY248). (**b-c**) Single-cells time-lapse quantification of transconjugants (**b**) bacterial and (**c**) nucleoid lengths. Average and SD are indicated (n cells analysed). The time of TAP acquisition (red dashed line at 0 min) corresponds to a 15 % increase in the green fluorescence in the transconjugant cells. (**d**) Upper panel: diagram of the fluorescent reporter system. Donor cells exhibit diffuse red fluorescence from the mCherry produced by TAP; recipients exhibit diffuse RecA-GFP fluorescence; transconjugants are identified by the production of red fluorescence followed by RecA-GFP polymerization. Lower panel: time-lapse microscopy images performed in a microfluidic chamber. Donor (D), recipient (R), and transconjugant (T) cells are indicated. Scale bar 1 μm. Donors TAP-Cas9-nsp (LY1537) or TAP-Cas9-lacZ2 (LY1538), recipient RecA-GFP (LY844). (**e-f**) Single-cells time lapse quantification of transconjugants (**e**) cell length and (**f**) skewness of RecA-GFP fluorescence signal. Average and SD are indicated (n cells analysed). The time of TAP acquisition (red dashed line at 0 min) corresponds to a 30 % increase in the green fluorescence in the transconjugant cells.

Using live-cell microscopy, we characterized the cellular response of the recipient cells to the acquisition of TAPs (Fig.2). In these experiments, the TAPs carry the sfGFP reporter system that confers green fluorescence to the donor and transconjugant cells. The lac+ recipient cells produce the nucleoid-association protein HU-mCherry, which localization reveals the global organization of the chromosome. As expected, the acquisition of the TAP-Cas9-nsp reported by the production of sfGFP green fluorescence in red recipient cells has no impact on growth, cell morphology or nucleoid organization (Fig.2a-c and movie 1). By contrast, the acquisition of the TAP-Cas9-lacZ2 triggers the rapid disorganization of the nucleoid that grows into an unstructured DNA bulk, followed by cells filamentation and occasional cell lysis (Fig.2a-c and movie 2). Furthermore, we analyzed in recipient cells the localization pattern of a RecA-GFP fusion, which has been reported to polymerize into large intracellular structures in response to DNA-damage induction^19^. In this experiment, TAPs carry the mCherry reporter system that confers red fluorescence to donors and transconjugant cells. Image analysis reveals that the acquisition of the TAP-Cas9-lacZ2 (Fig.2d and movie 3), but not the TAP-Cas9-nsp (Fig.2d and movie 4), is followed by cells filamentation (Fig.2e) as well as the RecA-GFP polymerization, which was quantified using fluorescence skewness analysis (Fig.2f, see methods). Nucleoid disorganization, cell filamentation and RecA-GFP bundle formation confirm that TAP-Cas9-lacZ2 acquisition is followed by CRISPR-mediated induction of DSBs that result in the death of the transconjugants.

### TAPs-mediated selective killing within a mixed E. coli recipient population

We verified the specificity of TAPs-mediated killing within a mixed recipient population composed of the targeted lac+ and the non-targeted lac- *E. coli* strains. We observe a ~4 log-fold decrease in viable lac+ transconjugants compared to lac-transconjugants when using the TAP-Cas9-lacZ2, while no difference is observed with the TAP-Cas9-nsp (Fig.3a). TAP-Cas9-lacZ2 specific killing activity is also reflected by the stagnation of the targeted lac+ recipient total population, while the non-targeted lac- population is able to grow during the six hours of mating (Fig.3b). We performed live-cell microscopy imaging where the lac+ and lac-recipients are distinguished by the typical localization pattern of nucleoid associated HU-mCherry and the replisome associated DNA clamp DnaN-mCherry, respectively. Time-lapse analysis shows that both strains receive the plasmids reported by the increase of green fluorescence, yet only the targeted lac+ transconjugants exhibits cell filamentation, symptomatic of Cas9-mediated DNA-damage induction (Fig.3c). These results recapitulate the effects obtained when using individual recipient strains, and demonstrate that the TAPs achieve selective killing of the targeted strain within a mixed population.

**Figure 3.**
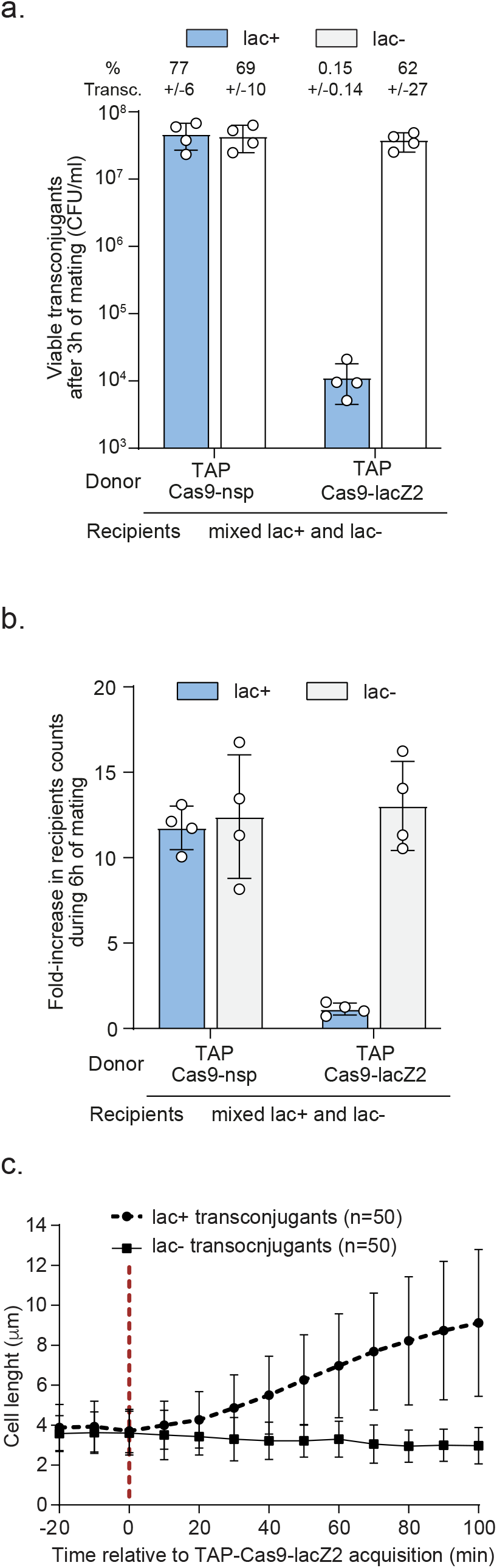
TAP system specifically kills targeted recipients in a mix of targeted and non-targeted *E. coli* recipient cells. (**a**) Viable transconjugant cells and percentage of transconjugants (ratio T/R+T) through TAP_kn_-Cas9-nsp or TAP_kn_-Cas9-lacZ2 transfer from donor to a mixed lac+ and lac-recipient population. (**b**) Quantification of fold-increase in lac+ and lac-recipient populations counts over the 6h of mating. Mean and SD are calculated from 4 independent experiments. Donors: TAP-Cas9-nsp (LY1369) or TAP-Cas9-lacZ2 (LY1370); recipients lac+ (LY827) and lac-(LY848). (**c**) Single-cell quantification showing cell length increase in the targeted lac+ transconjugant cells but not non-targeted lac-transconjugants. The time of TAP acquisition (red dashed line at 0 min) corresponds to a 15 % increase in the green fluorescence in the transconjugant cells. Cell length average is indicated with SD (n cells analysed). Donor TAP-Cas9-lacZ2 (LY1380); recipients HU-mCherry lac+ (LY248) and DnaN-mCherry lac-(LY1423).

### Analysis of TAP-escape mutants

Transfer of the TAP-Cas9-lacZ2 is associated with a ~3.5-log viability loss of the lac+ transconjugant cells, yet we noticed a proportion of transconjugants that are able to survive despite the acquisition of the TAP (Fig.1d). Genotyping and sequence analysis of 31 clones escaping the TAP-Cas9-lacZ2 activity revealed two types of escape mutants (Supplementary Fig.3a-b and supplementary discussion). One third (12 out of 31) have acquired a transposase or IS insertion in the plasmid-born *cas9* gene, thus inactivating the CRISPR system. Two-third have acquired mutations that modify the targeted *lacZ* chromosome locus, either by small or large deletions (12 out of 31) as already described^11^, or by single point mutation in the seed region of the PAM (7 out of 31), which was shown to be key for recognition by the Cas9-gRNA complex^20^ (Supplementary Fig.3c).

### TAPs directed against carbapenem-resistant population

Conjugative plasmids are major contributors to the spread of multi-drug resistance in bacteria^21^, those conferring carbapenem resistance being of severe clinical concern^22^. The IncL/M pOXA-48a plasmid carries the *blaOXA-48* gene that encodes the OXA-48 carbapenemase, which confer resistance to carbapenem and other beta lactams, such as imipenem and penicillin^23^. We designed TAPs targeting the pOXA-48a and assessed their ability to sensitize the plasmid-carrying population to ampicillin. Using an OXA48 spacer that targets the 5’-end of the *blaOXA*-*48* gene, we constructed the TAP-Cas9-OXA48 to induce Cas9-mediated DSBs on pOXA-48a, and the TAP-dCas9-OXA48 to inhibit *blaOXA*-*48* gene transcription by CRISPRi (Fig.4a). Transfer of TAP-Cas9-OXA48 and TAP-dCas9-OXA48 plasmids into pOXA-48a-carrying *E. coli* recipients lead to a ~4.5-log decrease in ampicillin resistance level, while the TAP-Cas9-nsp or the TAP-dCas9-nsp have no effect (Fig.4b). Unexpectedly, we observe that significantly less viable transconjugant are obtained without ampicillin selection when using the TAP-Cas9-OXA48 compared to TAP-dCas9-OXA48 (Fig.4b). We ruled out the possibility of a decrease in TAP-Cas9-OXA48 transfer ability as all four tested plasmids are acquired with similar frequency by pOXA-48a plasmid-free *E. coli* recipients (Supplementary Fig.4). However, analysis of the pOXA-48a plasmid sequence revealed the presence of the *pemIK* toxin-antitoxin (TA) system, which is involved in plasmid stability by inhibiting the growth of daughter cells that do not inherit the plasmid^24–26^. Indeed, the arrest of *pemIK* expression due to plasmid loss results in the rapid depletion of the labile PemI antitoxin, which can no longer repress the toxic activity of the more stable PemK toxin. This regulation was reported using CRISPR-associated phage therapy to cure antibiotic resistance carried by the pSHV-18 plasmid^26^. We then hypothesized that the observed reduction of viable TAP-Cas9-OXA48 transconjugants could be due to PemK toxic activity in cells that have lost of the pOXA-48a targeted by the Cas9 cleavage. This possibility was confirmed by inserting a constitutively expressed antitoxin *pemI* gene into the TAP-Cas9-OXA48, which results in a ~1.5 log increase in transconjugants viability, while retaining the inhibition of ampicillin resistance (Fig.4b).

**Figure 4.**
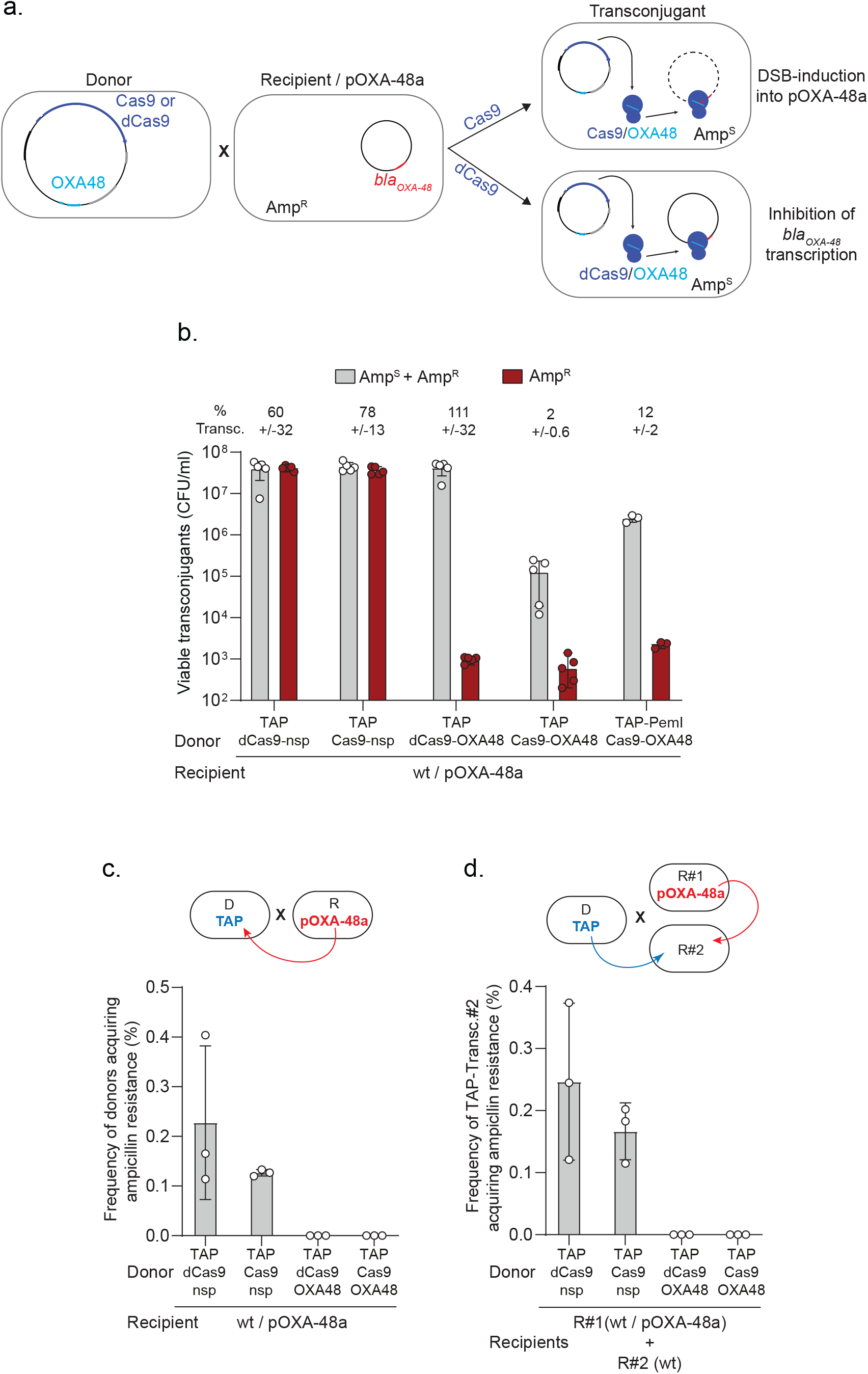
TAP re-sensitises pOXA48-carrying recipient cells and impedes resistance dissemination. (**a**) Diagram of the TAP-mediated anti-resistance strategy. TAP-Cas9-OXA48 targeting the *blaOXA48* promoter is transferred from a donor to an ampicillin resistant recipient cells carrying the pOXA-48a plasmid. Acquisition of the TAP-Cas9-OXA48 induces double-strand-breaks (DSBs) into the plasmid, while the TAP-dCas9-OXA48 inhibits *blaOXA48* gene expression. Both TAPs sensitise the transconjugant cells to ampicillin. (**b**) Histograms showing reduction of ampicillin resistance in transconjugants cells after acquisition of the TAP_kn_-Cas9-OXA48, TAP_kn_-dCas9-OXA48 and TAP_kn_-Cas9-OXA48-PemI. Percentages of transconjugants (ratio T/R+T) are indicated. (**c**) Histograms showing the frequency of donor cells acquiring ampicillin resistance through transfer of pOXA-48a from the recipients (as depicted in the above diagram). (**d**) Histograms show the frequency of ampicillin-resistance acquisition through pOXA-48a transfer into R#2 plasmid-free wt recipient that have received the TAPs (TAP-transc.#2) (as depicted in the above diagram). Mean and SD are calculated from at least 3 independent experiments. Donors TAP-dCas9-nsp (LY1524), TAP-Cas9-nsp (LY1369), TAP-dCas9-OXA48 (LY1523), TAP-Cas9-OXA48 (LY1522) or TAP-PemI-Cas9-OXA48 (LY1549); Recipients R#1 wt / pOXA-48a (LY1507) and R#2 wt (LY945).

The pOXA-48a is an autonomous conjugative plasmid that disseminates among *Enterobacteriaceae*, raising the possibility that the recipient containing the pOXA-48a could transfer ampicillin resistance to the TAPs-donors during mating. We observed that ampicillin resistance is indeed transmitted to ~0.2 % and 0.12 % of donors carrying the TAP-dCas9-nsp or the TAP-Cas9-nsp (Fig.4c). However, donors carrying the TAP-dCas9-OXA48 or the TAP-Cas9-OXA48 do not acquire ampicillin resistance (Fig.4c). Assuming that the efficiency of pOXA48 transfer is insensitive to the presence of the TAPs in the cells, this result suggests that TAPs directed against OXA48 impedes the development of resistance, even if the pOXA48 plasmid is acquired. We tested this possibility by performing the same conjugation experiments with an additional plasmid-free recipient wt strain (R#2) in the conjugation mix (see diagram in Fig.4d). Among R#2 cells that have received the TAP-Cas9-nsp or the TAP-dCas9-nsp, ~0.24 % and 0.15 % become ampicillin resistant, respectively. However, no ampicillin resistance is observed in R#2 cells that have received the TAP-Cas9-OXA48 or the TAP-dCas9-OXA48 (Fig.4d). Altogether, these results demonstrate that directing TAPs against the *blaOXA*-48 gene is an efficient strategy to sensitize the pOXA-48a-carrying strain to ampicillin. In addition, TAPs also appear to impede drug-resistance dissemination by protecting the donor and other plasmid-free recipients from developing the resistance.

### CSTB software: targeting specific strains within multispecies bacterial population

Designing TAPs that perform antibacterial activity against specific bacterial species, without affecting other bacterial strains, requires the robust identification of spacer sequences that are present in the genome of the targeted organism(s), but not in the genomes of other non-targeted strains. To complete this task, we developed a “Crispr Search Tool for Bacteria” CSTB algorithm, that enables the comparative analysis of ~18-23 nt long spacer sequences across a wide range of bacterial genomes and plasmids. The CSTB back-end database indexes all occurrences of these motifs present in 2919 complete genomes classified according to the NCBI taxonomy. CSTB allow identifying appropriate spacer sequences to reprogram TAPs against unique or multiple sites in the targeted chromosome or plasmid DNA.

We asked the CSTB algorithm to generate spacer sequences that target the attachment/effacement (A/E) pathogen *Citrobacter rodentium* strain ICC168 (Cr spacers), or the enteropathogenic *E. coli* EPEC strain E2348/69 (EPEC spacer), or the nosocomial pathogen *Enterobacter cloacae* (Ecl spacer), or the three of them (EEC spacer), without targeting any other bacterial genome present in the database. TAPs directed against *C. rodentium* carry a Cr1 spacer that target a unique locus, or a Cr22 that targets 22 loci distributed throughout the genome (Supplementary Fig.5a). Transfer of TAP-Cas9-Cr1 from an *E. coli* donor reduces by 4-log the viability of *C. rodentium* transconjugant cells (Supplementary Fig.5b). Live-cell microscopy revealed that TAP-Cas9-Cr1 acquisition induces *C. rodentium* filamentation and lysis, while no growth defect was induced by the TAP-Cas9-nsp (Supplementary Fig.5c and d; movies 5 and 6). This indicates that, as observed in *E. coli,* the induction of a single DSB by the Cas9 is lethal to *C. rodentium*. Consistently, targeting 22 chromosome loci by the TAP-Cas9-Cr22 results in comparable transconjugant killing efficiency (Supplementary Fig.5b and supplementary discussion). However, multiple targeting unbalances the contribution of the mechanisms by which transconjugants escape to the TAPs activity. Analysis of twenty clones escaping Cr1 single targeting revealed either deletions of the targeted chromosomal locus or inactivation of the CRISPR system on the TAP, in equal proportion. By contrast, the vast majority (19 out of 20) of clones escaping the Cr22 multiple targeting carry mutations that inactivate the TAP CRISPR system (Supplementary Fig.6 and Supplementary discussion). This is consistent with the prediction that mutations of the 22 targeted chromosome sites within the same call is highly infrequent, if even possible.

TAP transfer through F conjugation machinery is highly efficient towards MG1655 *E. coli* laboratory strain reaching up to 90% efficiency in 3h of mating (Fig.1, Fig.2 and Supplementary Fig.2). We quantified the efficiency of TAP-Cas9-nsp transfer in non-laboratory strain and observed a disparity between recipients with an overall ~7- to 900-fold decrease in TAP acquisition frequency in comparison to MG1655 *E. coli* (Fig.5a). To account for this variability, we normalized the frequency of viable transconjugants obtained for TAP-Cas9-Cr1, -Ecl, -EPEC and -EEC to the frequency of TAP-Cas9-nsp transconjugants in the corresponding bacterial strain (Fig.5b). We quantified that the TAP-Cas9-Cr1 induces a transconjugant viability loss only in *C. rodentium*, TAP-Cas9-Ecl in *E. cloacae*, TAP-Cas9-EPEC in *E. coli* EPEC, while the TAP-Cas9-EEC targets the three pathogenic strains. As a control, we show that the commensal *E. coli* HS recipient, which genome is not targeted by any spacer, is affected by none of these antibacterial TAPs (Fig.5b). These results demonstrate that the spacer sequences generated by the CSTB algorithm allow the robust reprograming of the TAPs for efficient and strain-specific antibacterial activity on mono-species recipient populations. It also demonstrates that one given TAP can target several species at the time.

**Figure 5.**
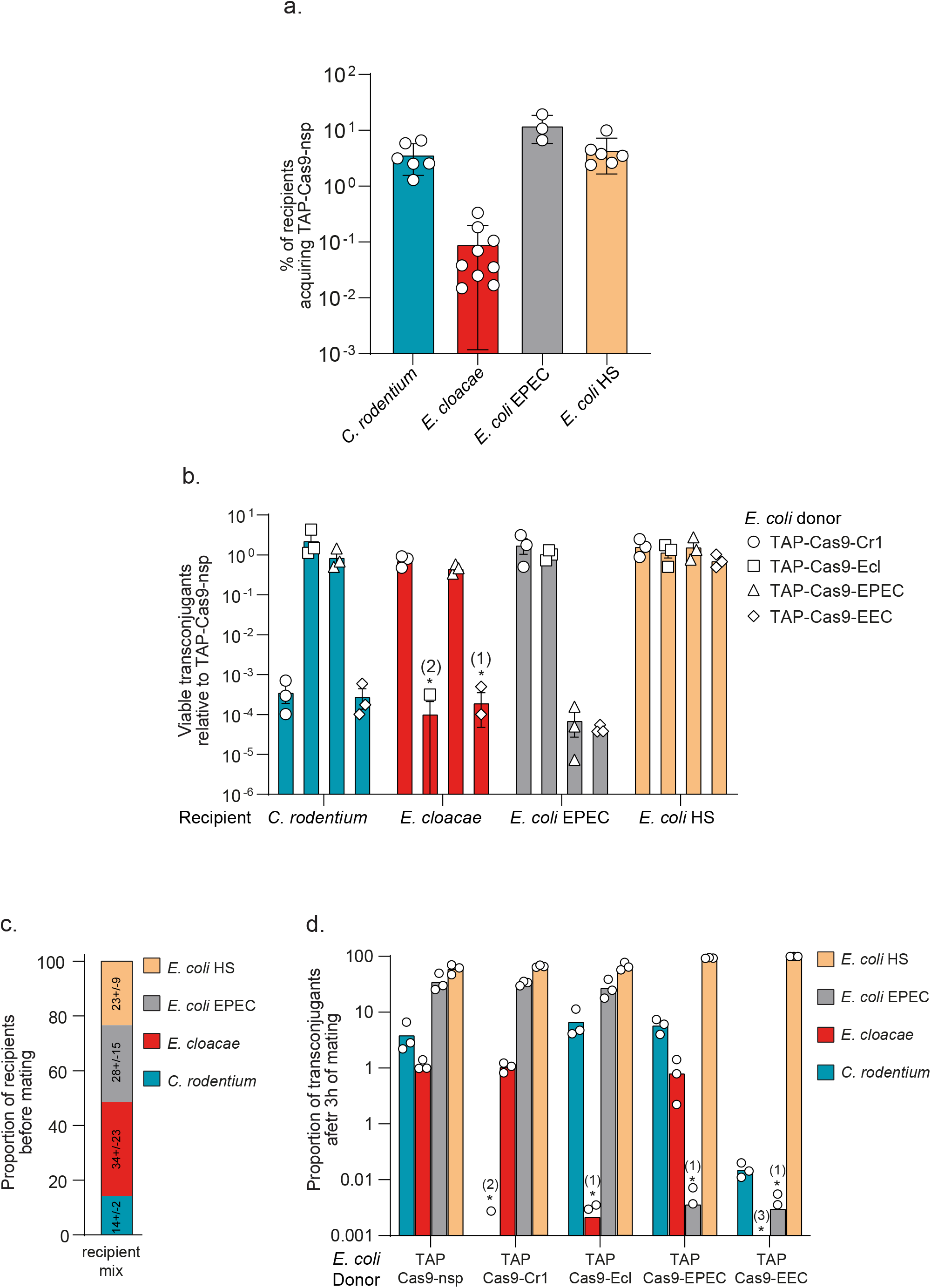
Efficient and strain-specific killing of TAPs within a multispecies recipient population. (**a**) Efficiency of TAP-Cas9-nsp transfer from *E. coli* (LY1369) donor to *C. rodentium*, *E. cloacae, E. coli* EPEC or HS recipients. Histograms show the percentages of tranconjugants (T/R+T) after 24h of conjugation for *C. rodentium*, *E. cloacae, E. coli* EPEC recipients and 3h for *E. coli* HS recipient; mean and SD are calculated from at least 3 independent experiments. (**b**) TAPs carrying specific spacers identified with the CSTB algorithm were tested against each recipient cells. To account for the variability of TAP transfer in the different recipient strains, the histograms show the relative abundance of viable transconjugants normalized by viable transconjugants obtained for the TAP_kn_-Cas9-nsp. Numbers in brackets above asterisk indicate replicates with detection limit of transconjugants below 10^−8^. Mean and SD are calculated from 3 independent experiments. (**c**) Proportion of recipients estimated by plating assay before mating with donors. Mean and SD calculated from 3 independent experiments are indicated for each recipient strains. (**d**) Each TAP carrying specific spacers were tested through conjugation between *E. coli* donors and a recipient population containing all recipient species. Histograms show the proportion of viable transconjugants in the mixed population after 3h of mating. Numbers in brackets above asterisk indicate replicates with detection limit of transconjugants below 10^−8^. Mean and SD are calculated from 3 independent experiments. Donors TAP-Cas9-nsp (LY1369), TAP-Cas9-Cr1 (LY1597), TAP-Cas9-Ecl (LY1566), TAP-Cas9-EPEC (LY1618), TAP-Cas9-EEC (LY1665); recipients *C. rodentium* (LY720), *E. cloacae* (LY1410), *E. coli* EPEC (LY1615) or HS (LY1601).

Next, we addressed TAPs ability to induce strain-specific antibacterial activity within a multi-species recipient population composed of an equal proportion the three pathogenic strains and the commensal *E. coli* HS (Fig.5c). The proportion of transconjugants obtained after 3h of mating with TAP-Cas9-nsp varies (Fig.5d) and reflects the efficiency of TAP transfer among the different recipient strains (Fig.5a). We observed that within the multispecies recipient mix, *C. rodentium* transconjugant viability is dramatically reduced by TAP-Cas9-Cr1, that of *E. cloacae* by TAP-Cas9-Ecl and that of *E. coli* EPEC by TAP-Cas9-EPEC, while all three species are affected by the triple-targeting TAP-Cas9-EEC. The viability of transconjugants of the control commensal *E. coli* HS is not affected by any of the antibacterial TAPs (Fig.5d). These results validate that TAPs achieve selective killing within a multispecies mixed recipient population without affecting the non-targeted species. Although the antibacterial TAPs impact selectively the viability of the transconjugant populations, their activity is not significantly reflected by the total recipient counts of each species (Supplementary Fig.7), due to the limited efficiency of TAP transfer to the pathogenic recipient strains and the differential fitness of these strains in competition within the conjugation mix.

## Discussion

Tools for *in situ* microbiota manipulation are currently in their infancy. Here we demonstrate the ability of the TAP antibacterial strategy to exert an efficient and strain-specific antibacterial activity within multi-species populations *in vitro*. TAPs selective-killing activity induces a ~4-log viability loss of the tested species. TAPs targeting the pOXA48a carbapenem resistance-plasmids results in a 4- to 5-logs increase of the strain susceptibility to the drug. Most CRISPR delivery methodology currently in development focus on the use of bacteriophages, which have intrinsically narrow host-range^27^. Besides, several recent studies successfully use the broad host range RK2 conjugation systems to deliver CRISPR system that target *E. coli*^26,28–30^ or *S. enterica*^31^ *in vitro*. One key advantage of our strategy over these approaches is the versatility conferred by the CSTB algorithm that enables the robust identification of gRNA that should be used to specifically re-target the TAPs against one or several bacterial strains of interest, without targeting other species. Despite the availability of numerous programs dedicated to the identification of CRISPR motifs, the CSTB has no equivalent so far^32^. TAPs can rapidly and easily be reprogrammed by changing the spacer sequence in one-step-cloning (see methods). Another advantage of TAPs is the constitutive expression of the CRISPR system (and the fluorescent reporters) from promoter that are active in a wide range of Enterobacteriaceae. The absence of requirement for an external inductor renders the TAP approach more suitable for the modification of natural bacterial communities *in vivo*.

Our work also reveals that TAPs efficiency is primarily determined by two main limiting factors. The first limiting factor is the ~10^−4^-10^−5^ frequency of escaper clones that acquire mutations inactivating the plasmid-born *cas9* gene, or mutations that modify the targeted sequence. As shown in *C. rodentium*, the latter escape mechanism can be avoided by targeting multiple sites on the genome of the targeted bacteria (see supplementary discussion). Another strategy would be to target essential genes, as their mutation is often lethal for the host bacteria^33^. The second limiting parameter is the efficiency of TAPs transfer towards the targeted strain(s). So far, all of the antibacterial^28,34^ or anti-drug^29,30,35^ methodology using conjugation are based on the incP RK2 conjugative system, which offer broad-host range, but low efficiency of transfer (10^−4^-10^−5^). Hamilton *et al*. have shown an artificial way to increase the efficiency of transfer using glass beads *in vitro* ^31^. Here, we use the F plasmid as a helper plasmid that mediates relatively efficient TAPs transfer (10^−1^-10^2^) to closely related Enterobacteriaceae. Therefore, TAPs appear appropriate to target a range of clinically relevant pathogenic or resistant bacteria (*E. coli*, *Citrobacter*, *Enterobacter*, *Klebsiella*, *Salmonella*, *Yersinia*, *Shigella*, *Serratia…*). Using TAPs to target phylogenetically distant species would require the development of broad-host range conjugation systems with increased transfer ability. Such superspreader plasmid mutants have been successfully isolated through Tn-seq approach^36,37^ and could represent an valuable option to widen the range of bacteria toward which TAPs could be directed.

Translating the present *in vitro* proof of concept to *in situ* settings would represent an important step towards the development of a non-antibiotic strategy for the *in situ* manipulation of microbiota composition, in a directed manner. TAPs could be used for the inhibition of harmful pathogenic and resistance strains from an infected host or environments, or as anti-virulence strategy through inhibition of virulence effector genes or genes involved in biofilm formation. The future applicability of the TAPs approach in clinical or environmental settings would require the consideration of the rapidly evolving regulations on GMO, CRISPR and biocontainment^38–40^.

## Methods

### Bacterial strains, plasmids, primer and growth culture conditions

#### Bacterial strains construction and growth procedures

Bacterial strains, plasmids and primers are listed in Tables S1, S2 and S3 respectively. Plasmid cloning were done by Gibson Assembly (Gibson *et al.*, 2009) and verified by Sanger sequencing (Eurofins Genomics). Chromosome mutation were transferred by phage P1 transduction to generate the final strains. Strains were grown at 37°C in Luria-Bertani (LB) broth, M9 medium supplemented with glucose (0.2%) and casamino acid (0.4%) (M9-CASA) or M63 medium supplemented with glucose (0.2%) and casamino acid (0.4%) (M63). When appropriate, the media were supplemented with the following antibiotics: 50 μg/ml kanamycin (Kan), 20 μg/ml chloramphenicol (Cm), 10 μg/ml tetracycline (Tc), 20 μg/ml nalidixic acid (Nal), 20 μg/ml streptomycin (St), 100 μg/ml ampicillin (Ap), 10 μg/ml gentamycin (Gm), 50 μg/ml rifampicin (Rif). When appropriate 40 μg/ml 5-bromo-4-chloro-3-indolyl-β-d-galactopyranoside (X-Gal) and 40 μM isopropyl β-D-1-thiogalactopyranoside (IPTG) were added for screening of LAC phenotype.

#### TAPs construction and one-step-cloning change of the spacer sequence on the TAPs

Plasmid construction was performed by IVA cloning (García-Nafría *et al.*, 2016), expect for changing the spacer sequence in the TAPs, which was performed by the replacement of the spacer in pEGL129 by a SapI-spacer-SapI DNA sequence. The nsp (non-specific) spacer sequence is flanked by two SapI restriction sites that allow for liberation of non-cohesive DNA ends upon SapI digestion. To replace the nsp spacer, a new spacer is constructed by annealing 2 oligonucleotides (listed in table S3) with complementary sequences to the non-cohesive ends generated by SapI restriction of TAP-Cas9-nsp or TAP-dCas9-nsp plasmids. Ligation production between the new spacer fragment and the TAP backbone was transformed into DH5α or TB28 strains. Constructions were verified by PCR reaction and sequencing.

### Congo red assay

#### Curli production colony assay for

*E. coli* strain OmpR234 with or without plasmids were plated on Congo Red medium (10 g bacto tryptone, 5 g yeast extract, 18 g bacto agar, 40 μg/ml Congo Red and 20 μg/ml Coomassie Brilliant blue G) and incubated 4 days at 30°C. Colonies were visualized at ×10 magnification with a M80 stereomicroscope (Leica). Digital images were captured with an IC80-HD integrated camera coupled to the stereomicroscope, operated via LASv4.8 software (Leica).

#### Liquid aggregation test

Overnight culture of *E. coli* strain OmpR234 with or without plasmids were diluted to an A_600_ of 0.05 in 1 ml M9-CASA medium supplemented with 25 μg/ml of Congo Red. Culture were grown without agitation at 30°C for 24h and image captured.

### Conjugation assay

Overnight cultures grown in LB of donor and recipient strains were diluted to an A_600_ of 0.05 and grown until an A_600_ comprised between 0.7 and 0.9 was reached. 50 μl of donor and 150μl of recipient cultures were mixed into an Eppendorf tube to obtain a 1:3 donor to recipient ratio. At time 0 min, 100 μl of the mix were diluted into 1 ml LB, serial diluted and plated on LB agar supplemented with antibiotics selecting for donor, recipient and transconjugant cells. The remaining 100 μl were incubated for 1h30 at 37°C. 1 ml of LB was added gently and the tubes were incubated at 37°C for another 1h30, 4h30 or 22h30. Conjugation mix were then vortexed, serial diluted and plated as for time 0 min.

#### Multispecies conjugation

Overnight cultures grown in LB of donor and recipient strains were diluted to an A_600_ of 0.05 and grown until an A_600_ comprised between 0.7 and 0.9 was reached. A recipient mix is prepared by mixing *C. rodentium*, *E. cloacae*, *E. coli* EPEC and *E. coli* HS recipients strains in equal proportion. This mix is serial diluted and plated on LB agar supplemented with antibiotics to select for each recipient. 100 μl of donor and 100 μl of the recipient mix were added to an Eppendorf tube to perform mating. At time 0 min, 100 μl of the mix were diluted into 1 ml LB, serial diluted and plated on LB agar supplemented with antibiotics to select for donor, recipients and transconjugants. The remaining 100 μl were incubated for 1h30 at 37°C. 1 ml of LB was gently added and the tubes were incubated for an additional 1h30 at 37°C. Conjugation mix were then vortexed, serial diluted and plated on LB agar supplemented with antibiotics to select for donor, recipients and transconjugants. In the figures, the efficiencies of conjugation are represented either as the final concentration of transconjugant cell (CFU/ml) or as the percentage of transconjugant cells calculated from the ratio (T/R+T).

### Transformation assay

Overnight cultures grown in LB were 1/100 diluted and grown until an A_600_ comprised between 0.4 and 0.6. Cells were treated with Rubidium Chloride and 90 μl of the resulting competent cells transformed with 100 ng of plasmid and heat shock. Following the 1h incubation at 37°C for phenotypic expression, cells were centrifugated 5 min at 5000 rpm, resuspended in 100 μl of LB, and 10 μl of serial dilution were spotted on LB plates supplemented with the appropriate antibiotics.

### Live-cell microscopy imaging and analysis

#### Time-lapse experiments

Overnight cultures in M9-CASA (between *E. coli*) or M63 (between *E. coli* and *C. rodentium*) of donor and recipient cells were diluted to an A_600_ of 0.05 and grown until an A_600_ comprised between 0.7 and 0.9. 25μl of donor and 75 μl of recipient were mixed into an Eppendorf tube and 50 μl of the mix was loaded into a B04A microfluidic chamber (ONIX, CellASIC^®^). Nutrient supply was maintained at 1 psi and the temperature maintained at 37°C throughout the imaging process. Cells were imaged every 10 min for 3 h.

#### Image acquisition

Conventional wide-field fluorescence microscopy imaging was carried out on an Eclipse Ti-E microscope (Nikon), equipped with x100/1.45 oil Plan Apo Lambda phase objective, FLash4 V2 CMOS camera (Hamamatsu), and using NIS software for image acquisition. Acquisition were performed using 50% power of a Fluo LED Spectra X light source at 488 nm and 560 nm excitation wavelengths. Exposure settings were 50 ms for sfGFP and 50 ms for mCherry produced from the TAPs; 100 ms for RecA-GFP; 100 ms for HU-mCherry; 100 ms for DnaN-mCherry.

#### Image analysis

Quantitative image analysis was done using Fiji software with MicrobeJ plugin (Ducret *et al.*, 2016). The Manual-editing interface of MicrobeJ was used to optimize cell detection and the Mean intensity fluorescence, skewness and cell length parameters were automatically extracted and plotted. We defined the timing of TAP acquisition (time t=0) by analyzing the increase of the fluorescence signal conferred by the TAPs (sfGFP or mCh). Plasmid acquisition was validated when a 15% sfGFP or a 30% mCherry fluorescence increase was observed in the transconjugant cells. Fluorescence profiles of each cells were then aligned according the defined t=0 to generate the graphs presented in Figures 2b, 2c, 2e, 2f, 3c and supplementary Fig. 5d.

### Flow cytometry

Conjugation was done as described in the conjugation assay section in 0.1 μm filtered LB. At time 90 min and 180 min, conjugation mix were diluted to an A_600_ of 0.03 in 0.1 μm filtered LB and analysed into an Attune NxT acoustic focusing cytometer at a 25 μl/min flow rate. Forward scattered (FSC), Side scattered (SSC) as well as fluorescence signal BL1 (sfGFP) and YL2 (mCherry) were acquired with the appropriate PMT setting and represented with the Attune™ NxT analysis software. To verify the absence of toxicity of the Cas9 or dCas9 constitutive expression from the TAPs, we compared the growth of *E. coli* MS388 / TAP with the *cas9* or *dcas9* or without any *cas9* gene. Those strains were grown overnight in 0.1 μm filtered LB and diluted to an A_600_ of 0.05 in 0.1 μm filtered LB. They were grown during 8h and the A_600_ and CFU/mL were estimated by plating assays at 0, 2, 4, 6 and 8 hours. In parallel, at 1h, 2h and 5h30 the strains were analysed into the Attune NxT acoustic Focusing cytometer at a 25 μl/min flow rate. Forward scattered (FSC) was acquired and represented with the Attune™ NxT software.

### Analysis of TAP-escape mutants

#### In E. coli

The 31 TAP-escape mutants were streaked on medium supplemented with X-Gal and IPTG to determine their LAC phenotype. TAP-escape mutants exhibiting lac+ phenotype were classified as “Blue” and the others as “White” in Supplementary Fig.3. To determine the acquisition of point mutation or deletion that modify the targeted lacZ2 locus, a PCR was realized with OL240 and OL654 that amplify a fragment of 748pb encompassing the lacZ2 locus in wt strain. For escape mutants that exhibited no deletion of the lacZ2 locus but still had an active TAP CRISPR system, the PCR product was sequenced and the mutations identified. A PCR was also done with OL655 and OL656 to amplify a larger fragment around lacZ2 and observe large deletion as previously described (Cui and Bikard, 2016). To determine the activity of the TAPs extracted from escape mutants, conjugation was performed between the TAP-escape mutants and an *E. coli* MS388 lac+ strain as described in the conjugation assays section. In parallel, the activity of the TAPs extracted with the Machery Nagel NucleoSpin^®^ Plasmid kit from escape mutants were verified by transformation into lac^+^ and lac^−^ strains as described in the transformation assay section. Seven inactive TAPs were sequenced to identify mutations inactivating CRISPR system.

#### C. rodentium

For the 20 TAP-Cas9-Cr1-escape mutants, a PCR with OL686 and OL687 was performed to determine deletions in the chromosome locus. To verify the CRISPR activity of the TAPs from *C. rodentium* TAP-escape mutants, conjugation was performed during 5 hours between the *C. rodentium* mutants and the *E. coli* MS388 strain to generate new *E. coli* TAPs donors. Then conjugation was performed during 24h between those new donors and fresh *C. rodentium* recipients and plated to select for recipient and transconjugants. To confirm the activity of the TAPs isolated form *C. rodentium* escape mutants, TAPs were extracted with Machery Nagel NucleoSpin^®^ Plasmid kit and transformed by electroporation (2,5kV) into wt *C. rodentium* cells treated with 10% sucrose. Following 1h of incubation at 37°C, cells were plated on LB-agar supplemented or not with Kan to evaluate the transformation efficiency. Two inactive TAP-Cas9-Cr1 and two inactive TAP-Cas9-Cr22 isolated from escaper clones were sequenced.

### CSTB algorithm

The CSTB web service enables the comparative analysis of CRISPR motifs across a wide range of bacterial genomes and plasmids. Currently considered motifs are NGG-anchored sequences of 18 to 23 base pairs long. The CSTB back-end database indexes all occurrences of these CRISPR motifs present in 2914 complete genomes labeled as representative or reference in the release 99 of RefSeq (03/12/20). In addition, 7 bacterial genomes and 5 plasmids of interest were added. The mean number of distinct motifs among bacterial genomes is 55923 (5719 and 2729570 as respective minimum and maximum). Genomes are classified according to the NCBI taxonomy (07/22/20). Each genome is inserted in the database of motifs by processing the corresponding complete fasta using the following procedure. Firstly, all words satisfying the CRISPR motif regular expression are detected and their chromosomal coordinates stored in a database of motifs. Secondly, all unique words are converted into an integer representation using a 2-bits per base encoding software we developed [https://github.com/glaunay/crispr-set]. These integers are then sorted in a unique flat file per genome. The indexing of CRISPR motifs as integers enables computationally efficient comparison of the sets of motifs across several organisms. Finally, the original fasta file is added to a blast database. All related software can be freely accessed at https://github.com/MMSB-MOBI/CSTB_database_manager. The CSTB input interface displays the 2914 genomes available for searching as two taxonomic trees. The left-hand tree allows for the selection of species whose genomes have to feature identical/similar CRISPR motifs. This set of genomes defines the targeted CRISPR motifs. Meanwhile, the right-hand tree allows for the selection of “excluded” organisms, which must have no motif in common with the targeted ones. The set of motifs that satisfies the user selections will effectively be equal to the union of the motifs found in the selected organisms subtracted from the intersection of the motifs found in the “excluded” organisms. Computation time ranges from seconds to a few minutes according to the size of the selections and an email is sent upon completion.

All the solutions CRISPR motifs are presented in an interactive table of sgRNA sequences and their occurrences in each selected organism. The table has sorting and filtering capabilities on motif counts and sequence composition. This allows for the easy selection of motifs of interest. Detailed information can be downloaded for the entire set of solutions or for the selected motifs only. This detailed information is provided in tabulated file with lines featuring the coordinates of each sgRNA motif in the targeted organisms. Alternatively, the user may explore the results using a genome-based approach. Hence, each targeted genome has its graphical view. The graphic is a circular histogram of the entire distribution of solution sgRNA motifs in a selected genome. The graphic is interactive to display the local breakdown of sgRNA distribution. The CSTB web site can be freely accessed at https://cstb.ibcp.fr.

## Acknowledgments

The authors thank Gregory Jubelin, François Cornet, Claire Prigent-Combaret, Pierre Bogaerts for the gift of strains and plasmids. Lisa Rubio, Brice Simon-Letcher, ShangNong Hu and Timothée Sluys, for earlier involvement in the project.

## Author contributions

C.L. provided funding. C.L and S.B. conceived the project, designed the study and wrote the paper. A.R., A.D. and S.B. performed the experiments and analyzed the data. The CSTB algorithm was conceived by G.L, C.H and S. L. in discussion with E.G., S.B., and C.L. All the authors gave intellectual input throughout the project.

## Supplementary information

### Supplementary discussion. TAP-escape mutations

TAPs designed to induce Cas9-mediated double-stranded-breaks (DSBs) to the chromosome of the targeted bacteria trigger significant viability loss. However, we observe the emergence of targeted bacteria that have evolved mutations allowing them to survive despite the acquisition of the TAP. The frequency of these TAP-escape mutants varies from ~3×10^−4^ to 6×10^−5^ depending on the spacer and the recipient strains used (Fig.1a, 3a, 4b and 5b). The phenotypic and sequence analysis of *E. coli* and *C. rodentium* escape-mutants reveal two main mechanisms to escape TAP activity: (i) The first mechanism is the acquisition of insertions (of transposase or Insertion Sequences, IS) or single nucleotide deletions that inactivate the *cas9* gene carried by the TAP. These types of mutations allowing bacteria to escape CRISPR activity have been previously reported to occur with comparable frequency ^1–4^. It has also been reported that mutations or deletions within either tracRNA or crRNA sequences is another way to inactivate CRISPR systems ^1–5^. Yet, no such mutations were found in the *C. rodentium* TAP-escape mutants analysed, potentially due to their lesser frequency of occurrence. (ii) The second mechanism we have identified is the acquisition of point mutations or deletion that modify the targeted locus, thus impeding the recognition by the gRNA. These were also previously described ^1–3,6,7^. When using TAPs that target one chromosome locus in *E. coli* and *C. rodentium*, escape-mutations occur by *cas9*-inactivation (first mechanism) in 38.7 % and 45 %, and by modification of the targeted locus (second mechanism) in 61.3 % and 55 %, correspondingly.

We show that the contribution of escape mutations by modification of the chromosome sequence can be dramatically, if not completely reduced by directing the TAPs against multiple chromosome loci. Indeed, the probability of mutating multiple chromosome sites within the same bacterial cell is expected to decrease as the number of targeted sites increases. When using the Cr22 spacer that targets 22 loci of *C. rodentium* chromosome, the vast majority of escape-mutations (19 out of 20 tested) carry mutations that inactivate the TAP-born *cas9* gene, and only one *C. rodentium* escape mutant still carry an active TAP and escapes through a still unknown mechanism. Consequently, we observe a 2.9-fold decrease in the frequency of TAP-Cas9-Cr22 escapers (8.56×10^−5^) compare to TAP-Cas9-Cr1 (2.47×10^−4^) that targets one single chromosome locus. This decrease is consistent with our estimates of the relative contributions of each escaping mechanisms. Interestingly, it has been shown that another way to avoid the emergence of escape mutants through the modification of the chromosome is to target essential genes, which mutation is often lethal for the cell ^5^.

The observe frequency of escape mutations by *cas9* inactivation is likely related to the intrinsic rate of transposition estimated to oscillate between 10^−5^ to 10^−6^ in *E. coli* ^8^ and the rate of spontaneous mutations (10^−8^ and 10^−10^ per base pair and generation). In the case of TAPs, it is also possible that the induction of DSBs results in the increase of the mutation rates through the triggering of SOS-induced hypermutator phenotype. Cui and Bikard demonstrated that one way to improve CRISPR efficiency/gRNA targeting in *E. coli* host cell is to inhibit RecA activity, which is essential to DSB repair and to the induction of the SOS-response ^6^. This strategy in however not relevant in the case of TAPs strategy, which is intended to target recipient strains that have not been genetically modified.

We also addressed the possibility that part of the mutations in TAPs already emerge in the donor cells, thus resulting in the transfer of already inactive TAPs into the recipient target. This possibility is supported by the sequencing analysis of one *C. rodentium* escape mutant that have received TAP-Cas9-Cr1 form an *E. coli* donor. The sequencing revealed the insertion in *cas9* of *insAB* genes coding for transposase elements present in *E. coli* but absent from *C. rodentium* genomes. This result indicates that mutation leading to TAPs inactivation can occur in the donor cells prior to transfer, without excluding that they might also emerge in the targeted recipient after plasmid acquisition.

## Supplementary figure legends

**Supplementary Figure 1.**
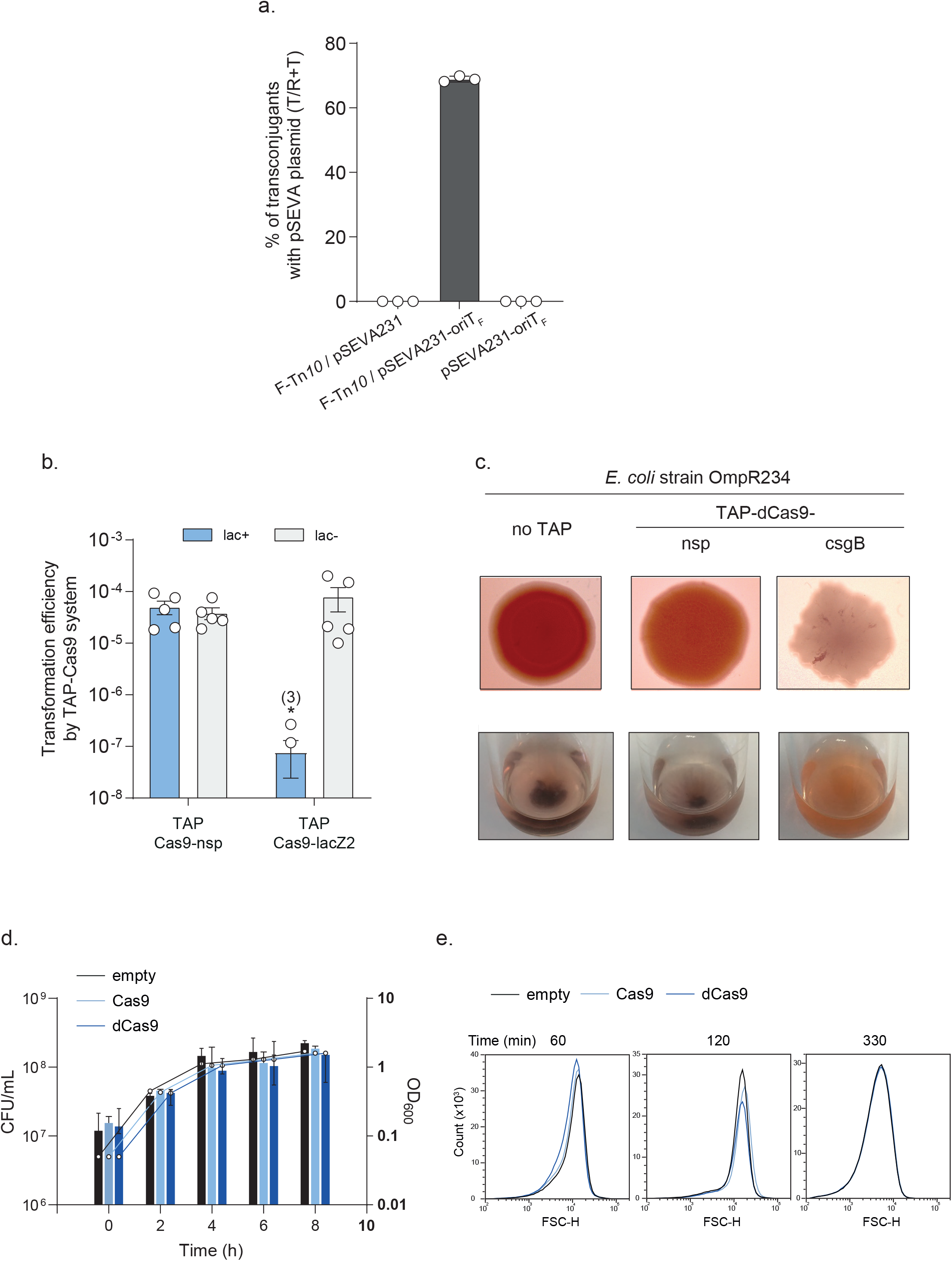
TAP carrying *oriT*_F_ are mobilized by the F plasmid Tra machinery and produces functional and non-toxic CRISPR and CRISPRi systems. (**a**) Plating assays showing the mobilization of the pSEVA plasmid carrying the F plasmid *oriT* by donors carrying the F-Tn*10* plasmid. Donors F-Tn*10*/pSEVA231 (LY887), F-Tn*10*/pSEVA231-oriTF (LY836), pSEVA231-oriT_F_ (LY874); recipient lac-(MS428). Mean and SD are calculated from 3 independent experiments. (**b**) Transformation efficiency of lac+ (MS388) or lac-(MS428) *E. coli* strains by TAP-Cas9-nsp or TAP-Cas9-lacZ2 (resistant to kn). Frequency of transformants is reported as the CFU numbers of kn resistant over total cells growing. Number in brackets above asterisk indicates replicates with transformant limit of detection below 10^−8^. Mean and SD are calculated from 5 independent experiments (**c**) Curli production analysis of OmpR234 *E. coli* strain carrying no TAP, TAP-dCas9-nsp or TAP-dCas9-csgB. Images of 5 days-old colony grown on congo red plates (top) and cell aggregats after overnight static growth in presence of congo red are shown. (**d-e**) Constitutive production of Cas9 or dCas9 from TAP has no effect on the *E. coli* growth (OD increase shown by lines) and viability (CFU/ml shown by histogram bars) (**d**) nor on the cell length analysed by flow cytometry (**e**). Mean and SD are calculated from 3 independent experiments. MS388 tested strain carries empty plasmid (pSEVA231-oriTF), Cas9 (TAP_kn_-Cas9-nsp) and dCas9 (TAP_kn_-dCas9-nsp).

**Supplementary Figure 2.**
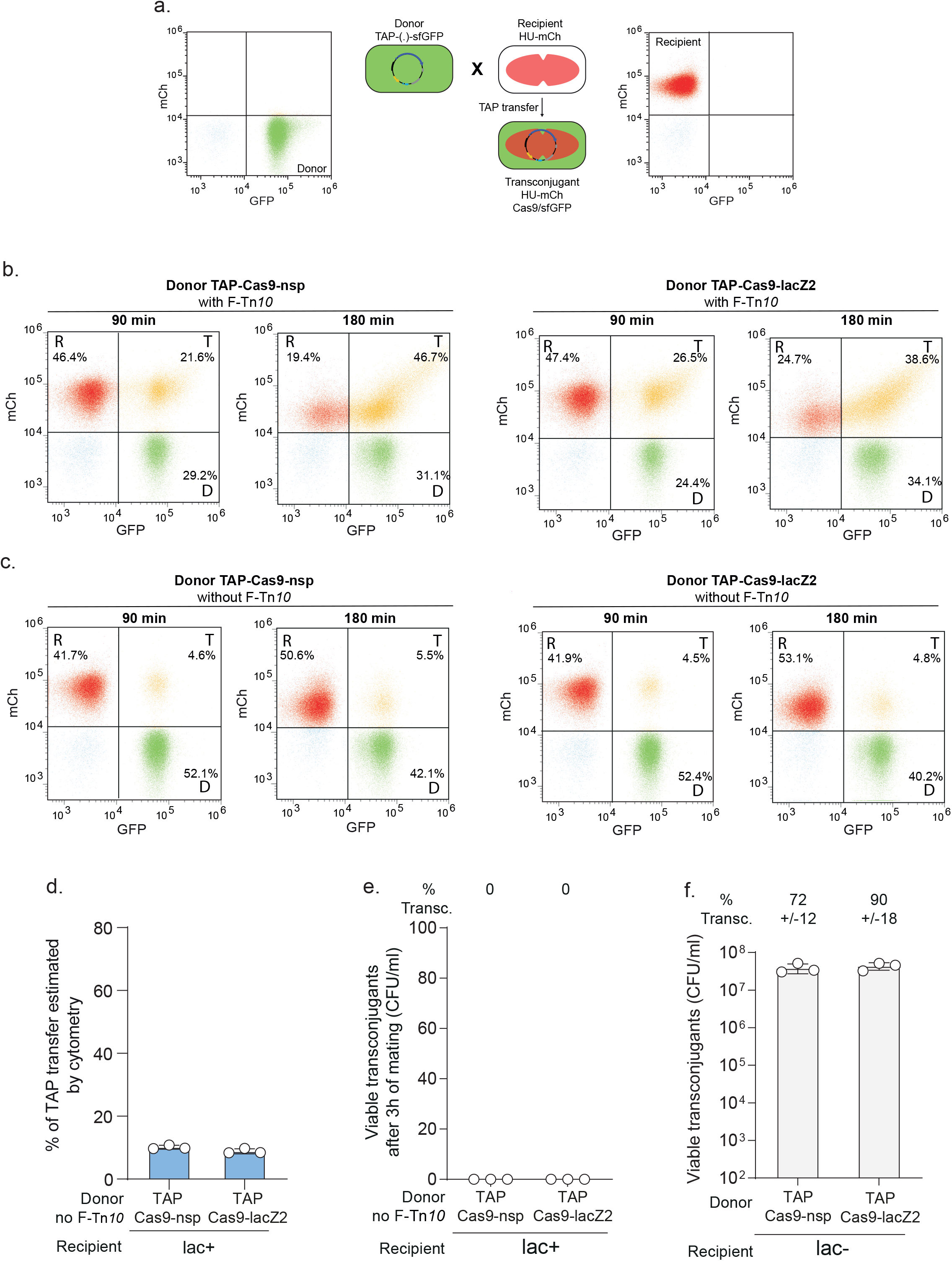
TAPs transfer equally in a F plasmid-dependent manner, regardless of the spacer sequence or the presence of the targeted chromosome locus in the recipient. (**a**) Diagram of the experimental system allowing the quantification of donor, recipient and transconjugant cells by flow cytometry. The TAP donor produces sfGFP green fluorescence (corresponding profile shown on the left panel). The recipient strains produce HU-mCherry red fluorescence from the chromosome (corresponding profile shown on the right panel). (**b**) The fraction of transconjugant cells producing both green and red fluorescence (yellow dots) increases similarly for TAP-Cas9-nsp and TAP-Cas9-lacZ2 after 90 and 180 min of mating when the donor contains the F plasmid. The proportion of each population is indicated in the corresponding region of interest (ROI) defined for donors (D), recipients (R) and transconjugant cells (T). (**c**) No transconjugant increase in observed with donor cells that do not carry the F-Tn*10* plasmid. Donors F-Tn*10* / TAP-Cas9-nsp (LY1371), F-Tn*10* / TAP-Cas9-lacZ2 (LY1380), TAP-Cas9-nsp without F-Tn*10* (LY1534), TAP-Cas9-lacZ2 without F-Tn*10* (LY1535); recipient lac+ (LY248). (**d**) A small percentage (~4-5 %) of particles exhibit both green and red fluorescence in the conjugation mixes using donor without F-Tn*10*. (**e**) The parallel plating of these conjugation mixes shows that they do not contain any transconjugant cells, indicating that these particles correspond to donor-recipient doublets. Donors TAP-Cas9-nsp without F-Tn*10* (LY1534), TAP-Cas9-lacZ2 without F-Tn*10* (LY1535); recipient lac+ (LY827). (**f**) Histogram showing the quantification of viable transconjugants cells obtained by transfer of the TAP-Cas9-nsp (LY1369) or TAP-Cas9-lacZ2 (LY1380) donors in lac-recipient cells (LY848) lacking the targeted sequence. Mean and SD are calculated from 3 independent experiments.

**Supplementary Figure 3.**
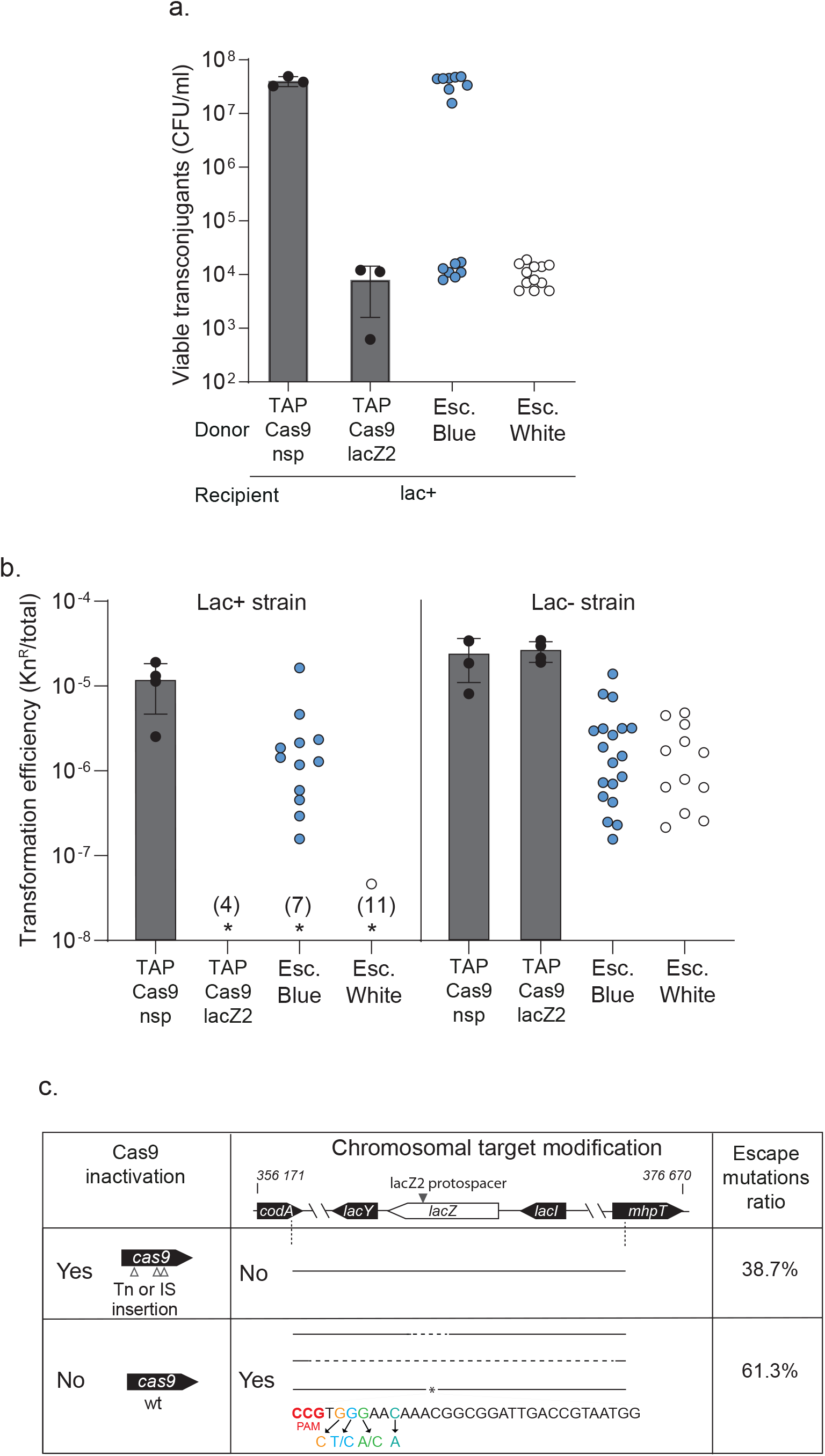
*E. coli* transconjugants escape to TAP by inactivating the TAP system or modifying the chromosomal target. (**a**) TAP-escape (esc.) mutants were distinguished according to their lac+ (blue) or lac-(white) phenotype on X-gal media reporting mutations in *lacZ*. TAPs functionality was analysed by conjugative assay mixing TAP-escape mutants as donors with lac+ recipient cells (LY832). Each dot represent viability of a single transconjugant replicate. Histograms report the mean and SD of viable transconjugants described in Fig.1d. (**b**) TAP-Cas9-nsp and TAP-Cas9-lacZ2 plasmids (carrying Kn resistance) extracted from white and blue escape mutants were transformed in lac+ (MS388) or lac-(MS428) strains. The transformation efficiency reported as ratio of Kn^R^ transformants over the total cell growing. Number in brackets above asterisk indicates replicates with transformant limit of detection below 10^−8^. (**c**) Summary of genotypic and sequence analysis of TAP-escape mutants. Schematic of the *lac* operon genomic region is shown with plain line representing no mutation while dotted lines and asterisk represent small or large deletions and single point mutation in the seed region of PAM respectively. Arrow indicates the position of the lacZ2 protospacer. Among TAP-escape analysed mutants, 38.7% (12 out of 31) have inactivated the TAP system and sequence analysis of 7 of them show transposase (Tn) or IS insertions in the *cas9* gene (InsH, InsAB from *E. coli* chromosome or IS*10* from F-Tn*10* plasmid). 61.3% of TAP-escape mutants (19 out 31) still carry an active TAP but modified the targeted sequence. Point mutations in seed region of 7 escape mutants does not affect the lac+ phenotype reported by their blue color on X-gal, thus explaining results shown in panel (a.) and (b.). 12 became lac-(white on X-gal) due to single nucleotide deletion in the seed region (1 out 12), 180 bp deletion within the *lacZ* gene (2 out 12) or larger deletion over 1 kb (9 out 12).

**Supplementary Figure 4.**
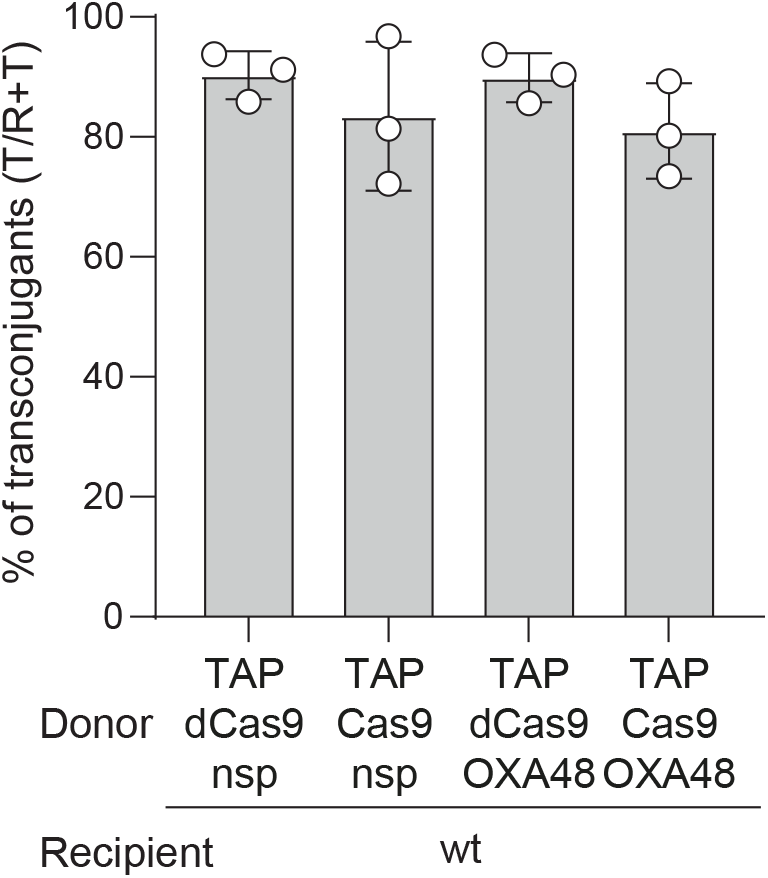
TAP-Cas9-nsp and TAP-Cas9-OXA48 targeting the *blaOXA48* promoter transfer with equal efficiency in pOXA-48a free recipient strain. Histograms of the percentage of transconjugants (T/R+T) recovered after mixing TAP-carrying donors with pOXA-48a free recipient cells. Mean and SD are calculated from 3 independent experiments. Donors TAP-dCas9-nsp (LY1524), TAP-Cas9-nsp (LY1369), TAP-dCas9-OXA48 (LY1523), or TAP-Cas9-OXA48 (LY1522); recipient wt (MS388).

**Supplementary Figure 5.**
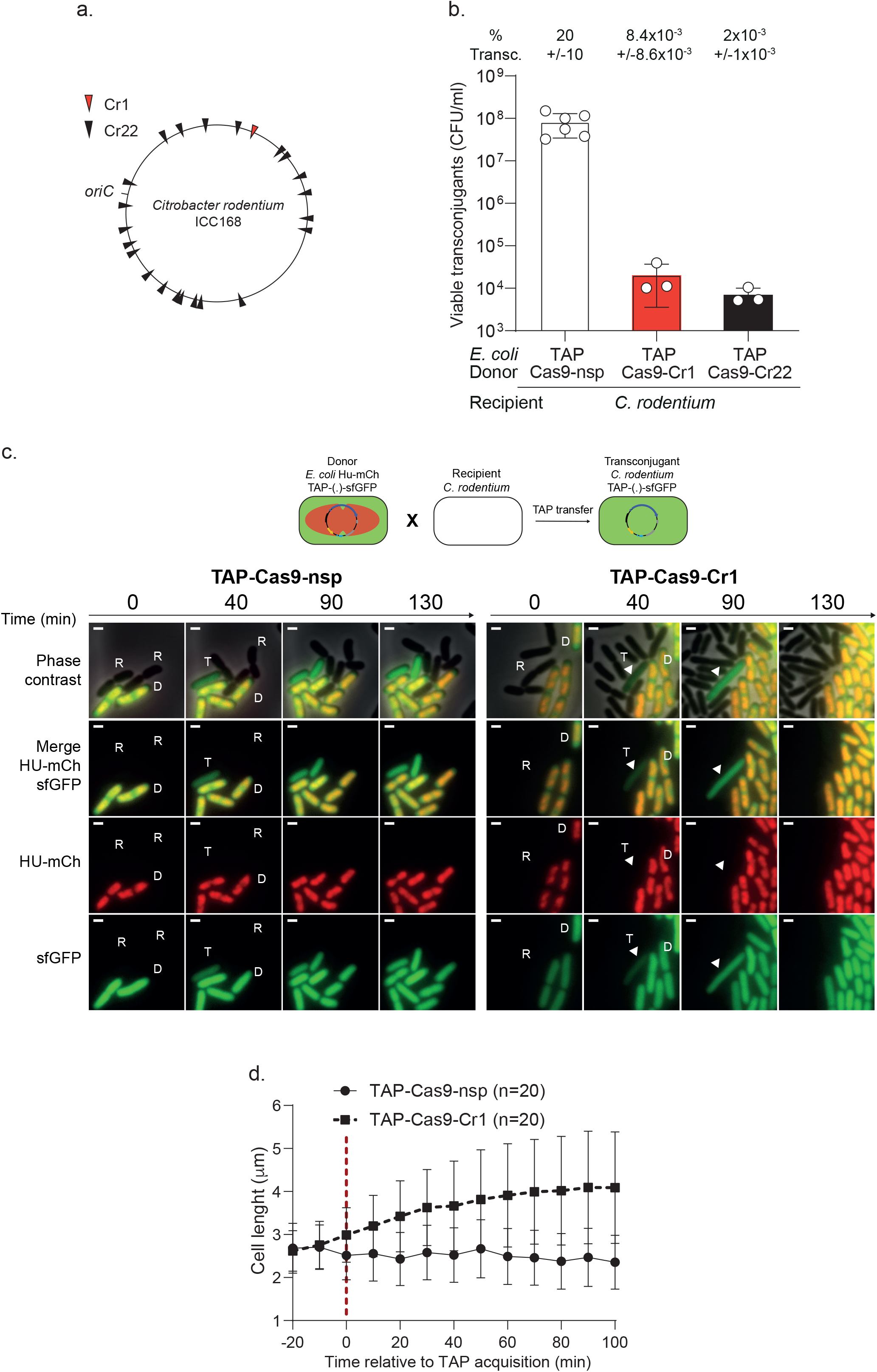
TAP mediates killing of *C. rodentium* by induction of a single double strand DNA break. (**a**) Map of the *C. rodentium* ICC168 chromosome with position of origin of replication (*oriC*) and the chromosome loci targeted by the Cr1 (red triangle) and Cr22 (black triangles) spacers. (**b**) Histogram of viable transconjugants through TAP transfer from *E. coli* donors to *C. rodentium* recipient cells. The percentage of transconjugants (T/R+T) obtained after 24h of mating are indicated above each bar. Mean and SD are calculated from at least 3 independent experiments. *E. coli* donors TAP-Cas9-nsp (LY1244), TAP-Cas9-Cr1 (LY1239) or TAP-Cas9-Cr22 (LY1276); recipient *C. rodentium* (LY720). (**c**) Visualization of TAP transfer from *E. coli* to *C. rodentium*. Top panel: diagram illustrating the system of transfer visualization. HU-mCherry *E. coli* donors carry sfGFP-producing TAP and exhibit both red and green fluorescence. *C. rodentium* recipients are not fluorescent and transconjugant cells exhibit green fluorescence after TAP acquisition. Lower panel: time-lapse microscopy images performed in a microfluidic chamber. D (donor), recipient (R), and transconjugant (T) cells are indicated. Scale bar 1 μm. Donors TAP-Cas9-nsp (LY1362) or TAP-Cas9-Cr1 (LY1363); recipient *C. rodentium* (LY720). (**d**) Single-cells time lapse quantification of bacterial length relative to the time of TAP acquisition (red dashed line at 0 min), identified by a 15 % increase in the green fluorescence in the transconjugant cells. Lines represent the population average with SD (n cells analysed).

**Supplementary Figure 6.**
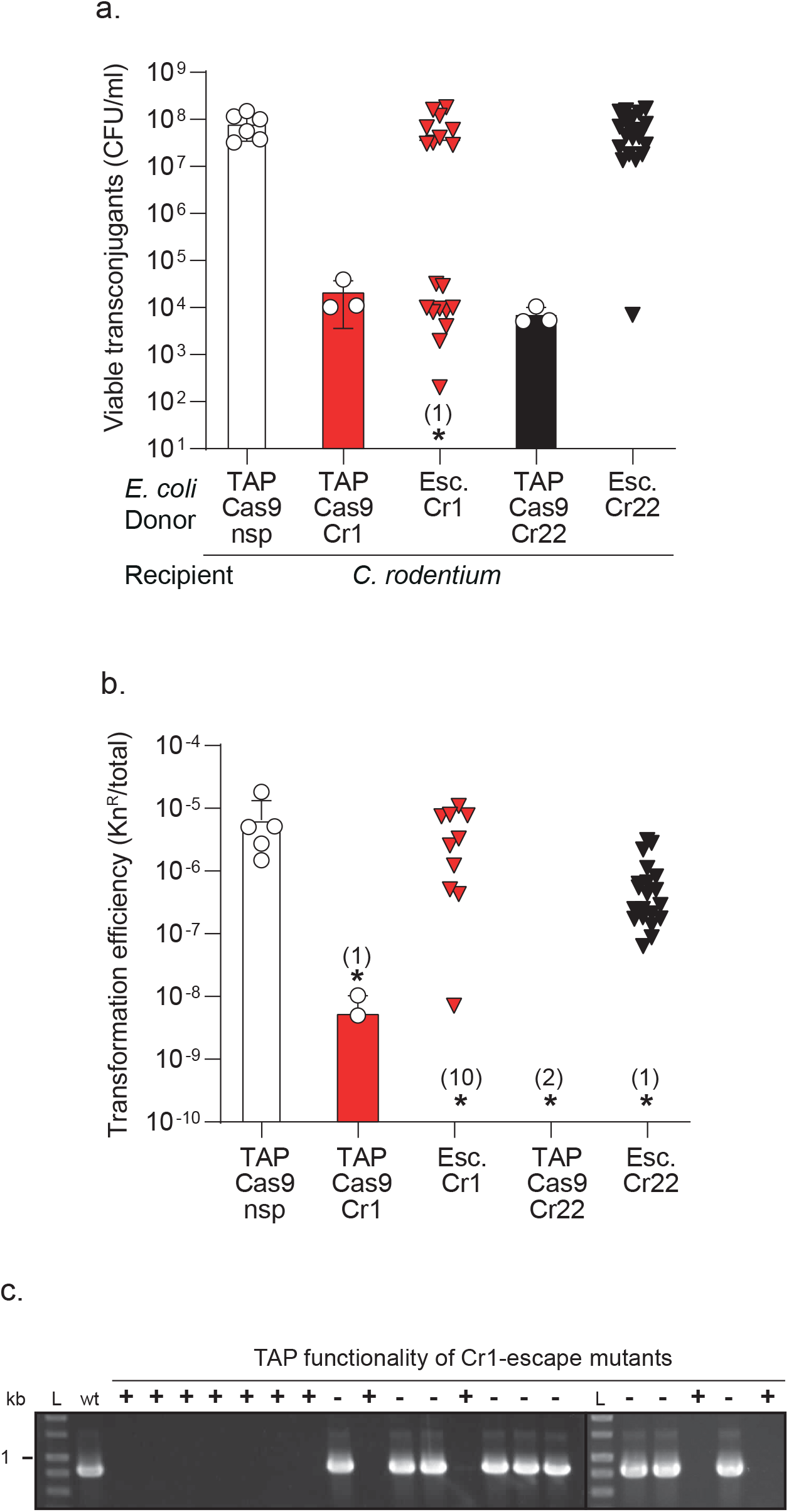
The mechanism of TAP-escape differs depending on number of targeted sites. **a**) TAPs activity analysed by mating assay between *E. coli* donors carrying TAPs isolated from *C. rodentium* escape mutants (Esc. Cr1 or Esc. Cr22), and *C. rodentium* recipients (LY720). Triangles represents the viability of each single transconjugant. Histograms report the mean and SD of viable transconjugants obtained in Supplementary Fig.5b. Number in brackets above asterisk indicate a replicate with a transconjugant limit of detection below 10^−7^. (**b**) Transformation efficiency of TAP-Cas9-nsp, TAP-Cas9-Cr1, TAP-Cas9-Cr22 and TAPs (carrying Kn resistance) extracted from *C. rodentium* escape mutants into *C. rodentium* (LY720). Frequency of transformants is reported as the CFU numbers of kn^R^ over total cells growing. Numbers in brackets above asterisk indicate replicates with transformant limit of detection below 10^−10^.(**c**) PCR analysis of the presence of the region encompassing the Cr1 protospacer region in *C. rodentium* mutants escaping the TAP-Cas9-Cr1 activity. Results show that the presence of inactive TAPs (labeled −) correlates with the presence of the intact target, while the presence of active TAP (labeled +) correlates with the deletion of the targeted sequence. L: ladder with kb scale.

**Supplementary Figure 7.**
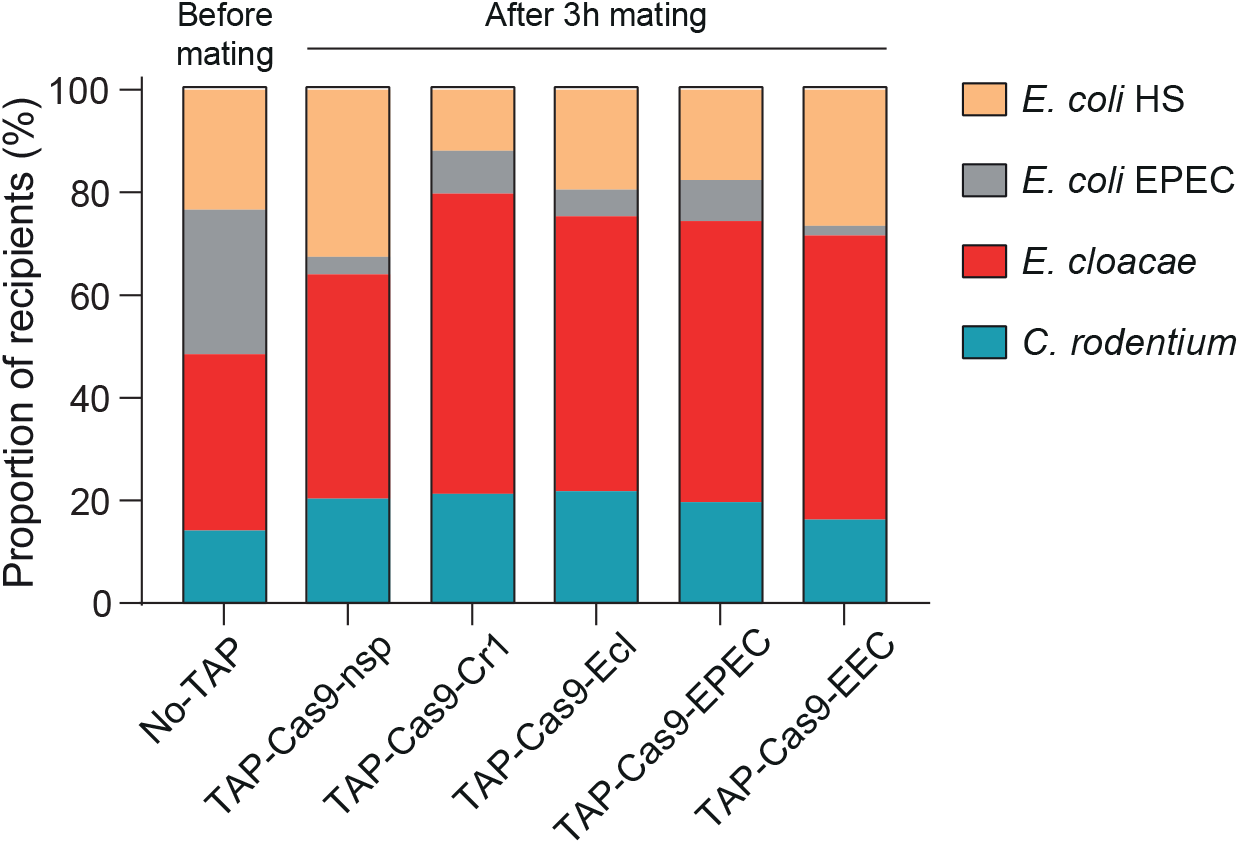
Targeted recipient total population is weakly impacted by transfer of TAPs in a multispecies conjugation mix. Histograms show the proportion of recipient cells estimated by plating assays after 3h of mating with TAP. Proportion of recipients before mating (No-TAP) reports the data shown in Fig.5c. Proportion mean are calculated from 3 independent experiments.

## Supplementary Movie legends

**Movie 1:** Microfluidic time-lapse imaging showing no impact of TAP-Cas9-nsp acquisition on nucleoid organization in *E. coli* (10 min per frame interval). Donors (LY1371) are revealed by green cytoplasmic sfGFP fluorescence, recipients (LY248) by nucleoid associated HU-mCherry fluorescence and transconjugant cells by combined HU-mCherry and sfGFP fluorescence. Left panel: merged phase contrast, sfGFP and Hu-mCherry fluorescence; middle panel: Hu-mCherry fluorescence; right panel: sfGFP fluorescence. Scale bar 1 μm.

**Movie 2:** Microfluidic time-lapse imaging showing the TAP-Cas9-lacZ2 acquisition followed by nucleoid decompaction and cell filamentation in *E. coli* (10 min per frame interval). Donors (LY1380) are revealed by green cytoplasmic sfGFP fluorescence, recipients (LY248) by nucleoid associated HU-mCherry fluorescence and transconjugant cells by combined HU-mCherry and sfGFP fluorescence. Left panel: merged phase contrast, sfGFP and Hu-mCherry fluorescence; middle panel: Hu-mCherry fluorescence; right panel: sfGFP fluorescence. Scale bar 1 μm.

**Movie 3:** Microfluidic time-lapse imaging showing no impact of TAP-Cas9-nsp acquisition on RecA-GFP dynamic in *E. coli* (10 min per frame interval). Donors (LY1537) are revealed by red cytoplasmic mCherry fluorescence, recipients (LY844) by diffuse RecA-GFP fluorescence and transconjugant cells by combined RecA-GFP and mCherry fluorescence. Left panel: merged phase contrast, RecA-GFP and mCherry fluorescence; middle panel: mCherry fluorescence; right panel: RecA-GFP fluorescence; middle panel: mCherry fluorescence. Scale bar 1 μm.

**Movie 4:** Microfluidic time-lapse imaging showing the TAP-Cas9-lacZ2 acquisition followed by RecA-GFP bundles formation and cell filamentation in *E. coli* (10 min per frame interval). Donors (LY1538) are revealed by red cytoplasmic mCherry fluorescence, recipient (LY844) cells by diffuse RecA-GFP fluorescence and transconjugant cells by the apparition mCherry fluorescence followed by RecA-GFP polymerization. Left panel: merged phase contrast, RecA-GFP and mCherry fluorescence; middle panel: mCherry fluorescence; right panel: RecA-GFP fluorescence; middle panel: mCherry fluorescence. Scale bar 1 μm.

**Movie 5:** Microfluidic time-lapse microscopy showing no impact of the TAP-Cas9-nsp acquisition on *C. rodentium* cell length (10 min per frame interval). Donors (LY1362) are revealed by nucleoid associated HU-mCherry and diffuse green cytoplasmic sfGFP fluorescence, *C. rodentium* wt recipient cells have no fluorescence and produce sfGFP fluorescence when they become transconjugants. Left panel: merged phase contrast, RecA-GFP and mCherry fluorescence; Middle panel: HU-mCherry fluorescence; Right panel: sfGFP fluorescence. Scale bar 1 μm.

**Movie 6:** Microfluidic time-lapse microscopy showing the TAP-Cas9-Cr1 acquisition by *C. rodentium* followed by cell length elongation (10 min per frame interval). Donors (LY1363) are revealed by nucleoid associated HU-mCherry and diffuse green cytoplasmic sfGFP fluorescence, *C. rodentium* wt recipient cells have no fluorescence and produce sfGFP fluorescence when they become transconjugants. Left panel: merged phase contrast, RecA-GFP and mCherry fluorescence; Middle panel: HU-mCherry fluorescence; Right panel: sfGFP fluorescence. Scale bar 1 μm.

## Supplementary Tables 1 to 3

**Supplementary Table 1:** strains.

| Name | Genotype | Reference |
| --- | --- | --- |
| LY720 | *Citrobacter rodentium* ICC168 (NaI<sup>R</sup>) | <sup>9</sup> |
| LY1602 | *Escherichia coli* EPEC E2348 | Gift from G. Jubelin |
| LY1615 | *Escherichia coli* EPEC E2348 (Rif<sup>R</sup>) | Derivative from LY1602 with a spontaneous mutation leading to rifampicin resistance phenotype |
| LY1563 | *Escherichia coli* HS | Gift from G. Jubelin |
| LY1601 | *Escherichia coli* HS (St<sup>R</sup>) | Derivative from LY1563 with a spontaneous mutation leading to streptomycin resistance phenotype |
| LY1410 | *Enterobacter cloacae* ATCC 13047 | Gift from P. Bogaerts |
| MS388 | *E. coli* MG1655 *rpsL* (St<sup>R</sup>, lac+) | Gift from F. Cornet |
| LY827 | MS388 *ilvA::Ap* (St<sup>R</sup>, Ap<sup>R</sup>, lac+) | <sup>10</sup> |
| MS428 | MS388 Δ*lacZ::gfp-parB<sub>PI</sub>* (St<sup>R</sup>, lac-) | Gift from F. Cornet |
| TB28 | *E. coli* K12 F- Lambda- *rph-I* DE(*lacIZYA*)::FRT (lac-) | Lab collection |
| DH5α | F<sup>-</sup> *endA1 glnV44 thi-1 recA1 relA1 gyrA96 deoR nupG purB20* ϕ80d*lacZ*ΔM15 $\Delta$(*lacZYA-argF*)U169, *hsdR17*(*r<sub>K</sub><sup>-</sup>m<sub>K</sub><sup>+</sup>*), $\lambda$ | Lab collection |
| OmpR234 | *E. coli* MG1655 *malA-kan ompR234* | Gift from C. Prigent-Combaret <sup>11</sup> |
| LY5 | MS388 / F-Tn10 (St<sup>R</sup>, Tc<sup>R</sup>, lac+) | Lab collection |
| LY6 | MS428 / F-Tn10 (St<sup>R</sup>, Tc<sup>R</sup>, lac-) | Lab collection |
| LY887 | MS388 / F-Tn10 / pSEVA231 (St<sup>R</sup>, Tc<sup>R</sup>, Kn<sup>R</sup>, lac-) | Transformation of LY5 with pSEVA231 |
| LY836 | MS388 / F-Tn10 / pBG6 (St<sup>R</sup>, Tc<sup>R</sup>, Kn<sup>R</sup>, lac-) | Transformation of LY5 with pBG6 |
| LY874 | MS388 / pBG6 (St<sup>R</sup>, Kn<sup>R</sup>, lac+) | Transformation of MS388 with pBG6 |
| LY636 | TB28 *ilvA::IsceICS-frt-cat-frt* (Cm<sup>R</sup>, lac-) | Lab collection |
| LY848 | MS428 *ilvA::erm* (St<sup>R</sup>, Erm<sup>R</sup>, lac-) | <sup>10</sup> |
| LY832 | MS388 *ilvA::erm* (St<sup>R</sup>, Erm<sup>R</sup>, lac+) | <sup>10</sup> |
| LY945 | MS388 *ilvA::cm* (St<sup>R</sup>, Cm<sup>R</sup>, lac+) | <sup>10</sup> |
| LY1361 | TB28 *ilvA::IsceICS-frt-cat-frt* / F-Tn10 (Cm<sup>R</sup>, Tc<sup>R</sup>, lac-) | Conjugation LY5 x LY636 to Cm<sup>R</sup> Tc<sup>R</sup> |
| LY1369 | TB28 *ilvA::IsceICS-frt-cat-frt* / F-Tn10 / pBG29 (Cm<sup>R</sup>, Tc<sup>R</sup>, Kn<sup>R</sup>, lac-) | Transformation of LY1361 with pBG29 |
| LY1370 | TB28 *ilvA::IsceICS-frt-cat-frt* / F-Tn10 / pAR5 (Cm<sup>R</sup>, Tc<sup>R</sup>, Kn<sup>R</sup>, lac-) | Transformation of LY1361 with pAR5 |
| LY1244 | MS388 / F-Tn10 / pBG29 (St<sup>R</sup>, Tc<sup>R</sup>, Kn<sup>R</sup>, lac+) | Transformation of LY5 with pBG29 |
| LY1239 | MS388 / F-Tn10 / pAR4 (St<sup>R</sup>, Tc<sup>R</sup>, Kn<sup>R</sup>, lac+) | Transformation of LY5 with pAR4 |
| LY1276 | MS388 / F-Tn10 / pAR7 (St<sup>R</sup>, Tc<sup>R</sup>, Kn<sup>R</sup>, lac+) | Transformation of LY5 with pAR7 |
| Ox468 | W1485 F<sup>-</sup> *leu thyA thi deoB* or *C supE rpsL, hupA-mCherry-kan* (St<sup>R</sup>, Kn<sup>R</sup>) | Gift from M. Stouf |
| LY119 | MS388 *hupA-mcherry-frt-Kan-frt* (St<sup>R</sup>, Kn<sup>R</sup>, lac+) | MS388 x P1 Ox468 to Kan<sup>R</sup> |
| LY248 | MS388 *hupA-mcherry-frt* (St<sup>R</sup>, lac+) | Derivative of LY119, *kan* removed via pCP20 |
| LY1056 | MS388 *hupA-mcherry-frt* / F-Tn10 (St<sup>R</sup>, Tc<sup>R</sup>, lac+) | Conjugation LY6 x LY248 to St<sup>R</sup> Tc<sup>R</sup> Kan<sup>R</sup> |
| LY844 | MG1655 *recA-gfp-kan, FhuB::recA-frt* | <sup>12</sup> |
| AB1157 | F<sup>-</sup>, λ-, *rac*-, *thi-1*, *hisG4*, $\Delta$(*gpt-proA*)62, *argE3*, *thr-1*, *leuB6*, *kdgK51*, *rfbC1*, *araC14*, *lacY1*, *galK2*, *xylA5*, *mtl-1*, *tsx-33*, *supE44*(*glnX44*), *rpoS396*, *rpsL31*(St<sup>R</sup>), *qsr'-0*, *mgl-51* | CGSC #1157 |
| LY183 | AB1157 *DnaN-mCherry-frt-kan-frt* (Kan<sup>R</sup>) | Gift from D. Sherratt <sup>13</sup> |
| LY1388 | MS428 *DnaN-mCherry-frt-kan-frt* (St<sup>R</sup>, Kan<sup>R</sup>, lac-) | P1 transduction from LY183 on MS428 |
| LY1423 | MS428 *DnaN-mCherry-frt* (St<sup>R</sup>, lac-) | Derivative of LY1388, *kan* removed via pCP20 |
| LY1371 | TB28 *ilvA::IsceICS-frt-cat-frt* / F-Tn10 / pAR1 (Cm<sup>R</sup>, Tc<sup>R</sup>, Kn<sup>R</sup>, lac-) | Transformation of LY1361 with pAR1 |
| LY1380 | TB28 *ilvA::IsceICS-frt-cat-frt* / F-Tn10 / pAR12 (Cm<sup>R</sup>, Tc<sup>R</sup>, Kn<sup>R</sup>, lac-) | Transformation of LY1361 with pAR12 |
| LY1362 | MS388 *hupA-mcherry-frt* / F-Tn10 / pAR1 (St<sup>R</sup>, Tc<sup>R</sup>, Kn<sup>R</sup>, lac+) | Transformation of LY1056 with pAR1 |
| LY1363 | MS388 *hupA-mcherry-frt* / F-Tn10 pAR11 (St<sup>R</sup>, Tc<sup>R</sup>, Kn<sup>R</sup>, lac+) | Transformation of LY1056 with pAR11 |
| LY1537 | TB28 *ilvA::IsceICS-frt-cat-frt* / F-Tn10 / pAR17 (Cm<sup>R</sup>, Tc<sup>R</sup>, Kn<sup>R</sup>, lac-) | Transformation of LY1361 with pAR17 |
| LY1538 | TB28 *ilvA::IsceICS-frt-cat-frt* / F-Tn10 / pAR18 (Cm<sup>R</sup>, Tc<sup>R</sup>, Kn<sup>R</sup>, lac-) | Transformation of LY1361 with pAR18 |
| LY1413 | *E. coli* Top10 / pOXA-48a (Ap<sup>R</sup>) | Gift from P. Bogaerts UCL Louvain |
| LY1507 | MS388 *ilvA::erm* / pOXA-48a (St<sup>R</sup>, Erm<sup>R</sup>, Ap<sup>R</sup>, lac+) | LY1413 x LY832 to Erm<sup>R</sup> Ap<sup>R</sup> |
| LY1522 | TB28 *ilvA::IsceICS-frt-cat-frt* / F-Tn10 / pBG50 (Cm<sup>R</sup>, Tc<sup>R</sup>, Kn<sup>R</sup>, lac-) | Transformation of LY1361 with pBG50 |
| LY1523 | TB28 *ilvA::IsceICS-frt-cat-frt* / F-Tn10 / pBG51 (Cm<sup>R</sup>, Tc<sup>R</sup>, Kn<sup>R</sup>, lac-) | Transformation of LY1361 with pBG51 |
| LY1524 | TB28 *ilvA::IsceICS-frt-cat-frt* / F-Tn10 / pAR6 (Cm<sup>R</sup>, Tc<sup>R</sup>, Kn<sup>R</sup>, lac-) | Transformation of LY1361 with pAR6 |
| LY1549 | TB28 *ilvA::IsceICS-frt-cat-frt* / F-Tn10 / pBG52 (Cm<sup>R</sup>, Tc<sup>R</sup>, Kn<sup>R</sup>, lac-) | Transformation of LY1361 with pBG52 |
| LY1618 | TB28 *ilvA::IsceICS-frt-cat-frt* / F-Tn10 / pAR28 (Cm<sup>R</sup>, Tc<sup>R</sup>, Kn<sup>R</sup>, lac-) | Transformation of LY1361 with pAR28 |
| LY1566 | TB28 *ilvA::IsceICS-frt-cat-frt* / F-Tn10 / pAR20 (Cm<sup>R</sup>, Tc<sup>R</sup>, Kn<sup>R</sup>, lac-) | Transformation of LY1361 with pAR20 |
| LY1665 | TB28 *ilvA::IsceICS-frt-cat-frt* / F-Tn10 / pAR29 (Cm<sup>R</sup>, Tc<sup>R</sup>, Kn<sup>R</sup>, lac-) | Transformation of LY1361 with pAR29 |
| LY1597 | TB28 *ilvA::IsceICS-frt-cat-frt* / F-Tn10 / pAR4 (Cm<sup>R</sup>, Tc<sup>R</sup>, Kn<sup>R</sup>, lac-) | Transformation of LY1361 with pAR4 |
| LY1534 | TB28 *ilvA::IsceICS-frt-cat-frt* / pBG29 (Cm<sup>R</sup>, Tc<sup>R</sup>, Kn<sup>R</sup>, lac-) | Transformation of LY636 with pBG29 |
| LY1535 | TB28 *ilvA::IsceICS-frt-cat-frt* / pAR5 (Cm<sup>R</sup>, Tc<sup>R</sup>, Kn<sup>R</sup>, lac-) | Transformation of LY636 with pAR5 |

**Supplementary Table 2:**
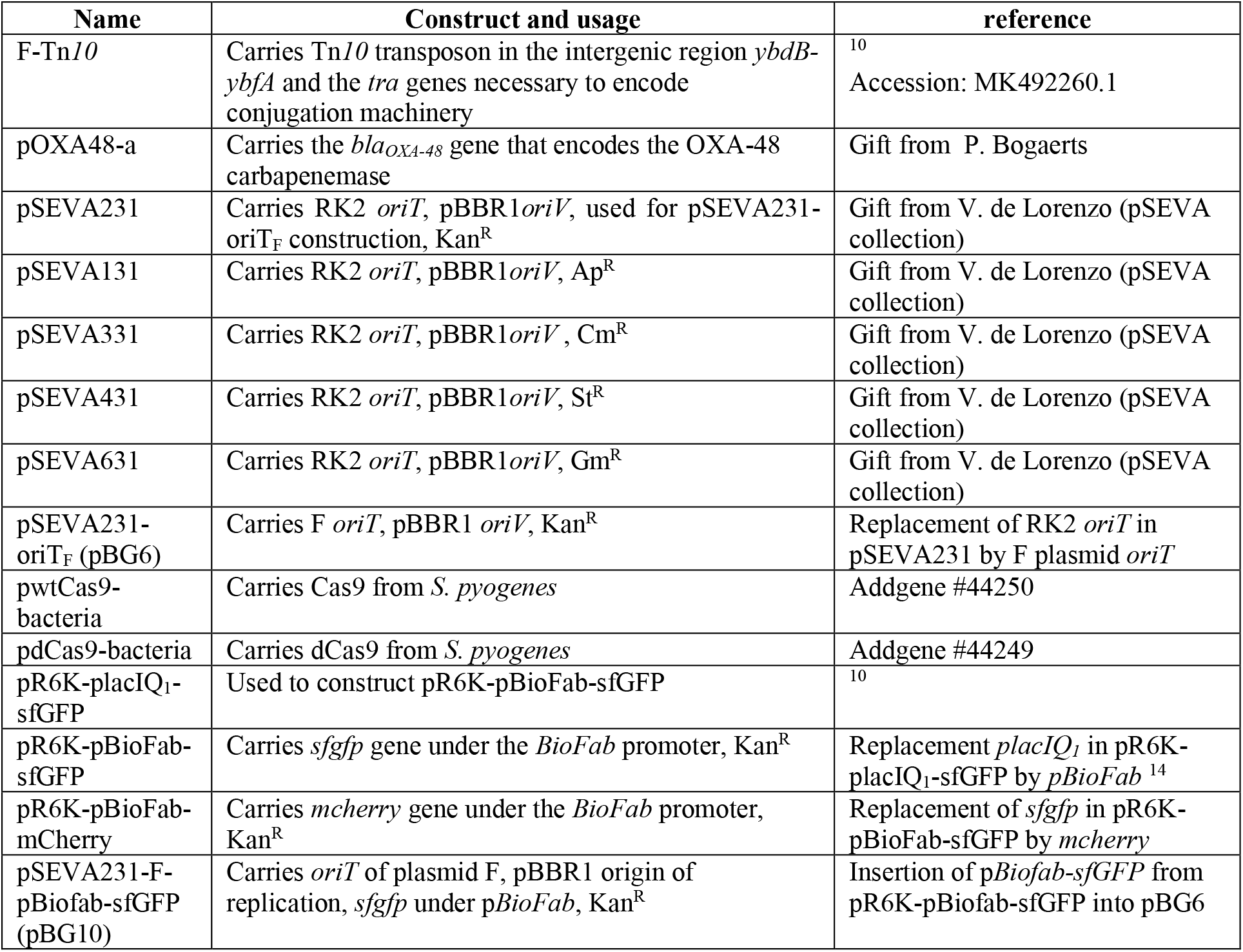
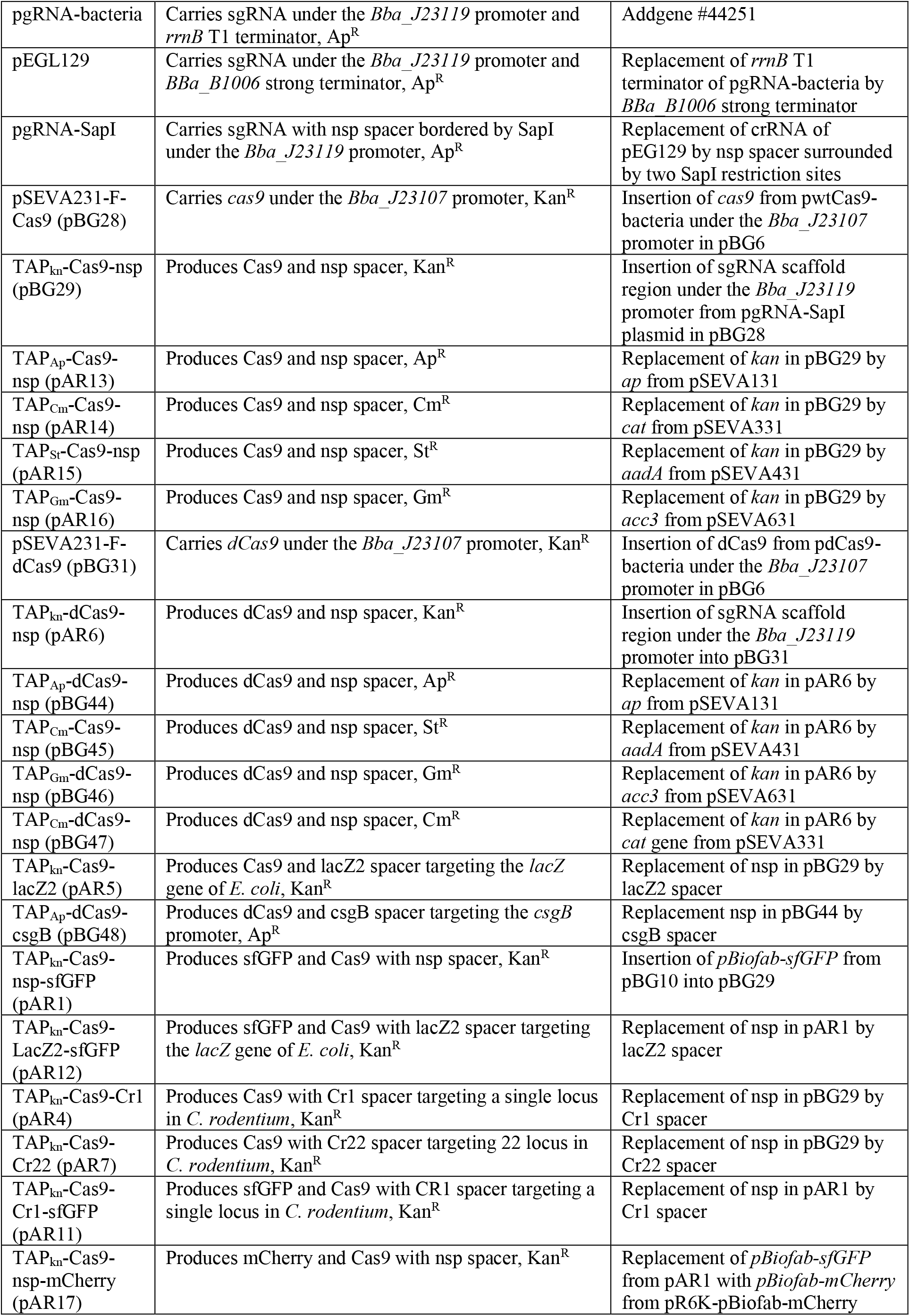
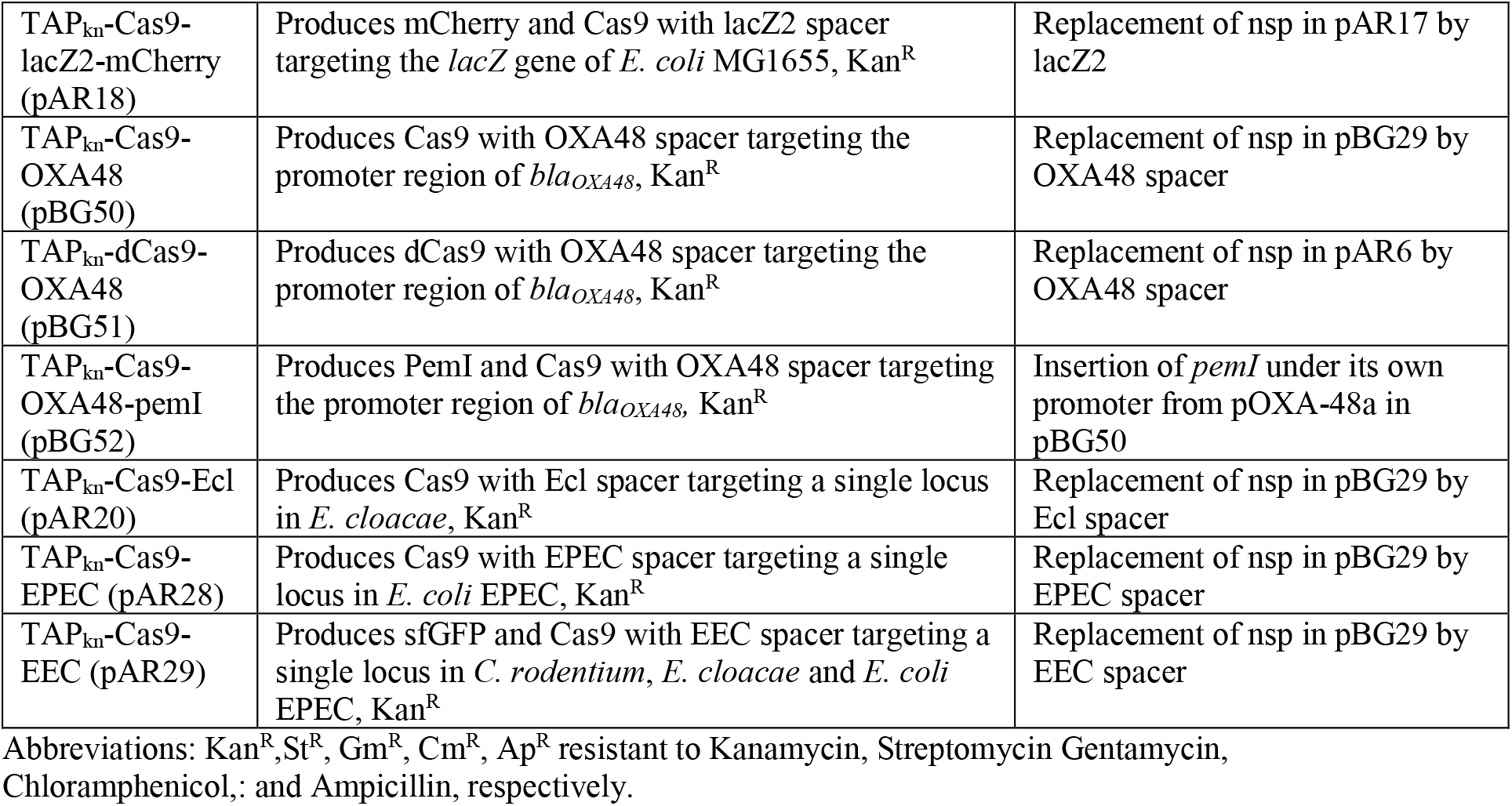
plasmids.

| Name | Construct and usage | reference |
| --- | --- | --- |
| F-Tn10 | Carries Tn10 transposon in the intergenic region *ybdB-ybfA* and the *tra* genes necessary to encode conjugation machinery | <sup>10</sup><br>Accession: MK492260.1 |
| pOXA48-a | Carries the *bla<sub>OXA-48</sub>* gene that encodes the OXA-48 carbapenemase | Gift from P. Bogaerts |
| pSEVA231 | Carries RK2 *oriT*, pBBR1*oriV*, used for pSEVA231-oriT<sub>F</sub> construction, Kan<sup>R</sup> | Gift from V. de Lorenzo (pSEVA collection) |
| pSEVA131 | Carries RK2 *oriT*, pBBR1*oriV*, Ap<sup>R</sup> | Gift from V. de Lorenzo (pSEVA collection) |
| pSEVA331 | Carries RK2 *oriT*, pBBR1*oriV*, Cm<sup>R</sup> | Gift from V. de Lorenzo (pSEVA collection) |
| pSEVA431 | Carries RK2 *oriT*, pBBR1*oriV*, St<sup>R</sup> | Gift from V. de Lorenzo (pSEVA collection) |
| pSEVA631 | Carries RK2 *oriT*, pBBR1*oriV*, Gm<sup>R</sup> | Gift from V. de Lorenzo (pSEVA collection) |
| pSEVA231-oriT<sub>F</sub> (pBG6) | Carries F *oriT*, pBBR1 *oriV*, Kan<sup>R</sup> | Replacement of RK2 *oriT* in pSEVA231 by F plasmid *oriT* |
| pwtCas9-bacteria | Carries Cas9 from *S. pyogenes* | Addgene #44250 |
| pdCas9-bacteria | Carries dCas9 from *S. pyogenes* | Addgene #44249 |
| pR6K-placIQ<sub>1</sub>-sfGFP | Used to construct pR6K-pBioFab-sfGFP | <sup>10</sup> |
| pR6K-pBioFab-sfGFP | Carries *sfgfp* gene under the *BioFab* promoter, Kan<sup>R</sup> | Replacement *placIQ<sub>1</sub>* in pR6K-placIQ<sub>1</sub>-sfGFP by *pBioFab* <sup>14</sup> |
| pR6K-pBioFab-mCherry | Carries *mcherry* gene under the *BioFab* promoter, Kan<sup>R</sup> | Replacement of *sfgfp* in pR6K-pBioFab-sfGFP by *mcherry* |
| pSEVA231-F-pBiofab-sfGFP (pBG10) | Carries *oriT* of plasmid F, pBBR1 origin of replication, *sfgfp* under *pBioFab*, Kan<sup>R</sup> | Insertion of *pBiofab-sfGFP* from pR6K-pBiofab-sfGFP into pBG6 |
| pgRNA-bacteria | Carries sgRNA under the *Bba_J23119* promoter and *rrnB* T1 terminator, Ap<sup>R</sup> | Addgene #44251 |
| pEGL129 | Carries sgRNA under the *Bba_J23119* promoter and *Bba_B1006* strong terminator, Ap<sup>R</sup> | Replacement of *rrnB* T1 terminator of pgRNA-bacteria by *Bba_B1006* strong terminator |
| pgRNA-SapI | Carries sgRNA with nsp spacer bordered by SapI under the *Bba_J23119* promoter, Ap<sup>R</sup> | Replacement of crRNA of pEG129 by nsp spacer surrounded by two SapI restriction sites |
| pSEVA231-F-Cas9 (pBG28) | Carries *cas9* under the *Bba_J23107* promoter, Kan<sup>R</sup> | Insertion of *cas9* from pwtCas9-bacteria under the *Bba_J23107* promoter in pBG6 |
| TAP<sub>kn</sub>-Cas9-nsp (pBG29) | Produces Cas9 and nsp spacer, Kan<sup>R</sup> | Insertion of sgRNA scaffold region under the *Bba_J23119* promoter from pgRNA-SapI plasmid in pBG28 |
| TAP<sub>Ap</sub>-Cas9-nsp (pAR13) | Produces Cas9 and nsp spacer, Ap<sup>R</sup> | Replacement of *kan* in pBG29 by *ap* from pSEVA131 |
| TAP<sub>Cm</sub>-Cas9-nsp (pAR14) | Produces Cas9 and nsp spacer, Cm<sup>R</sup> | Replacement of *kan* in pBG29 by *cat* from pSEVA331 |
| TAP<sub>St</sub>-Cas9-nsp (pAR15) | Produces Cas9 and nsp spacer, St<sup>R</sup> | Replacement of *kan* in pBG29 by *aadA* from pSEVA431 |
| TAP<sub>Gm</sub>-Cas9-nsp (pAR16) | Produces Cas9 and nsp spacer, Gm<sup>R</sup> | Replacement of *kan* in pBG29 by *acc3* from pSEVA631 |
| pSEVA231-F-dCas9 (pBG31) | Carries *dCas9* under the *Bba_J23107* promoter, Kan<sup>R</sup> | Insertion of dCas9 from pdCas9-bacteria under the *Bba_J23107* promoter in pBG6 |
| TAP<sub>kn</sub>-dCas9-nsp (pAR6) | Produces dCas9 and nsp spacer, Kan<sup>R</sup> | Insertion of sgRNA scaffold region under the *Bba_J23119* promoter into pBG31 |
| TAP<sub>Ap</sub>-dCas9-nsp (pBG44) | Produces dCas9 and nsp spacer, Ap<sup>R</sup> | Replacement of *kan* in pAR6 by *ap* from pSEVA131 |
| TAP<sub>Cm</sub>-dCas9-nsp (pBG45) | Produces dCas9 and nsp spacer, St<sup>R</sup> | Replacement of *kan* in pAR6 by *aadA* from pSEVA431 |
| TAP<sub>Gm</sub>-dCas9-nsp (pBG46) | Produces dCas9 and nsp spacer, Gm<sup>R</sup> | Replacement of *kan* in pAR6 by *acc3* from pSEVA631 |
| TAP<sub>Cm</sub>-dCas9-nsp (pBG47) | Produces dCas9 and nsp spacer, Cm<sup>R</sup> | Replacement of *kan* in pAR6 by *cat* gene from pSEVA331 |
| TAP<sub>kn</sub>-Cas9-lacZ2 (pAR5) | Produces Cas9 and lacZ2 spacer targeting the *lacZ* gene of *E. coli*, Kan<sup>R</sup> | Replacement of nsp in pBG29 by lacZ2 spacer |
| TAP<sub>Ap</sub>-dCas9-csgB (pBG48) | Produces dCas9 and csgB spacer targeting the *csgB* promoter, Ap<sup>R</sup> | Replacement nsp in pBG44 by csgB spacer |
| TAP<sub>kn</sub>-Cas9-nsp-sfGFP (pAR1) | Produces sfGFP and Cas9 with nsp spacer, Kan<sup>R</sup> | Insertion of *pBiofab-sfGFP* from pBG10 into pBG29 |
| TAP<sub>kn</sub>-Cas9-LacZ2-sfGFP (pAR12) | Produces sfGFP and Cas9 with lacZ2 spacer targeting the *lacZ* gene of *E. coli*, Kan<sup>R</sup> | Replacement of nsp in pAR1 by lacZ2 spacer |
| TAP<sub>kn</sub>-Cas9-Cr1 (pAR4) | Produces Cas9 with Cr1 spacer targeting a single locus in *C. rodentium*, Kan<sup>R</sup> | Replacement of nsp in pBG29 by Cr1 spacer |
| TAP<sub>kn</sub>-Cas9-Cr22 (pAR7) | Produces Cas9 with Cr22 spacer targeting 22 locus in *C. rodentium*, Kan<sup>R</sup> | Replacement of nsp in pBG29 by Cr22 spacer |
| TAP<sub>kn</sub>-Cas9-Cr1-sfGFP (pAR11) | Produces sfGFP and Cas9 with CR1 spacer targeting a single locus in *C. rodentium*, Kan<sup>R</sup> | Replacement of nsp in pAR1 by Cr1 spacer |
| TAP<sub>kn</sub>-Cas9-nsp-mCherry (pAR17) | Produces mCherry and Cas9 with nsp spacer, Kan<sup>R</sup> | Replacement of *pBiofab-sfGFP* from pAR1 with *pBiofab-mCherry* from pR6K-pBiofab-mCherry |
| TAP<sub>kn</sub>-Cas9-lacZ2-mCherry (pAR18) | Produces mCherry and Cas9 with lacZ2 spacer targeting the *lacZ* gene of *E. coli* MG1655, Kan<sup>R</sup> | Replacement of nsp in pAR17 by lacZ2 |
| TAP<sub>kn</sub>-Cas9-OXA48 (pBG50) | Produces Cas9 with OXA48 spacer targeting the promoter region of *bla*<sub>OXA48</sub>, Kan<sup>R</sup> | Replacement of nsp in pBG29 by OXA48 spacer |
| TAP<sub>kn</sub>-dCas9-OXA48 (pBG51) | Produces dCas9 with OXA48 spacer targeting the promoter region of *bla*<sub>OXA48</sub>, Kan<sup>R</sup> | Replacement of nsp in pAR6 by OXA48 spacer |
| TAP<sub>kn</sub>-Cas9-OXA48-pemI (pBG52) | Produces PemI and Cas9 with OXA48 spacer targeting the promoter region of *bla*<sub>OXA48</sub>, Kan<sup>R</sup> | Insertion of *pemI* under its own promoter from pOXA-48a in pBG50 |
| TAP<sub>kn</sub>-Cas9-Ecl (pAR20) | Produces Cas9 with Ecl spacer targeting a single locus in *E. cloacae*, Kan<sup>R</sup> | Replacement of nsp in pBG29 by Ecl spacer |
| TAP<sub>kn</sub>-Cas9-EPEC (pAR28) | Produces Cas9 with EPEC spacer targeting a single locus in *E. coli* EPEC, Kan<sup>R</sup> | Replacement of nsp in pBG29 by EPEC spacer |
| TAP<sub>kn</sub>-Cas9-EEC (pAR29) | Produces sfGFP and Cas9 with EEC spacer targeting a single locus in *C. rodentium*, *E. cloacae* and *E. coli* EPEC, Kan<sup>R</sup> | Replacement of nsp in pBG29 by EEC spacer |

Abbreviations: Kan<sup>R</sup>, St<sup>R</sup>, Gm<sup>R</sup>, Cm<sup>R</sup>, Ap<sup>R</sup> resistant to Kanamycin, Streptomycin Gentamycin, Chloramphenicol, and Ampicillin, respectively.

**Supplementary Table 3:**
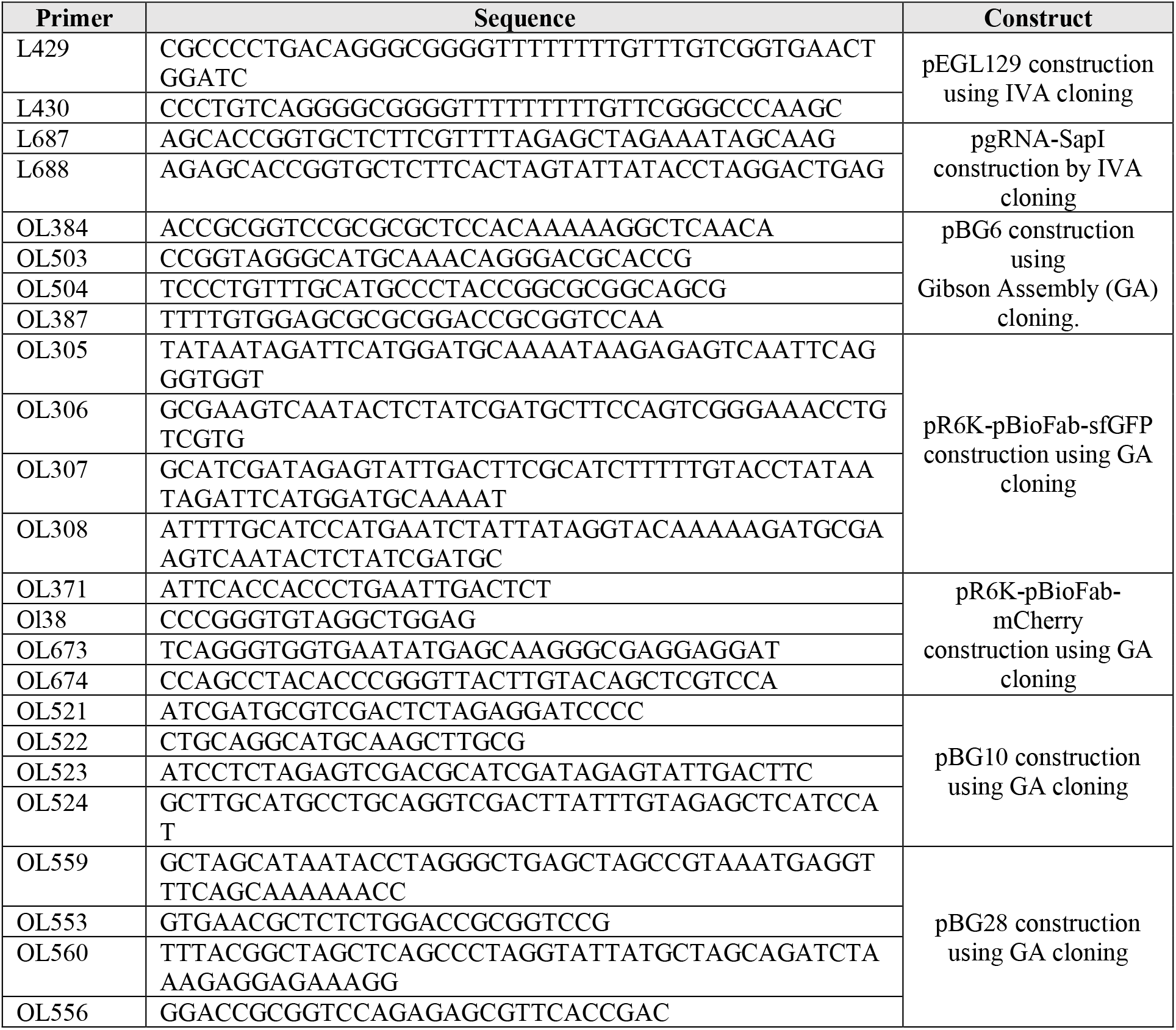
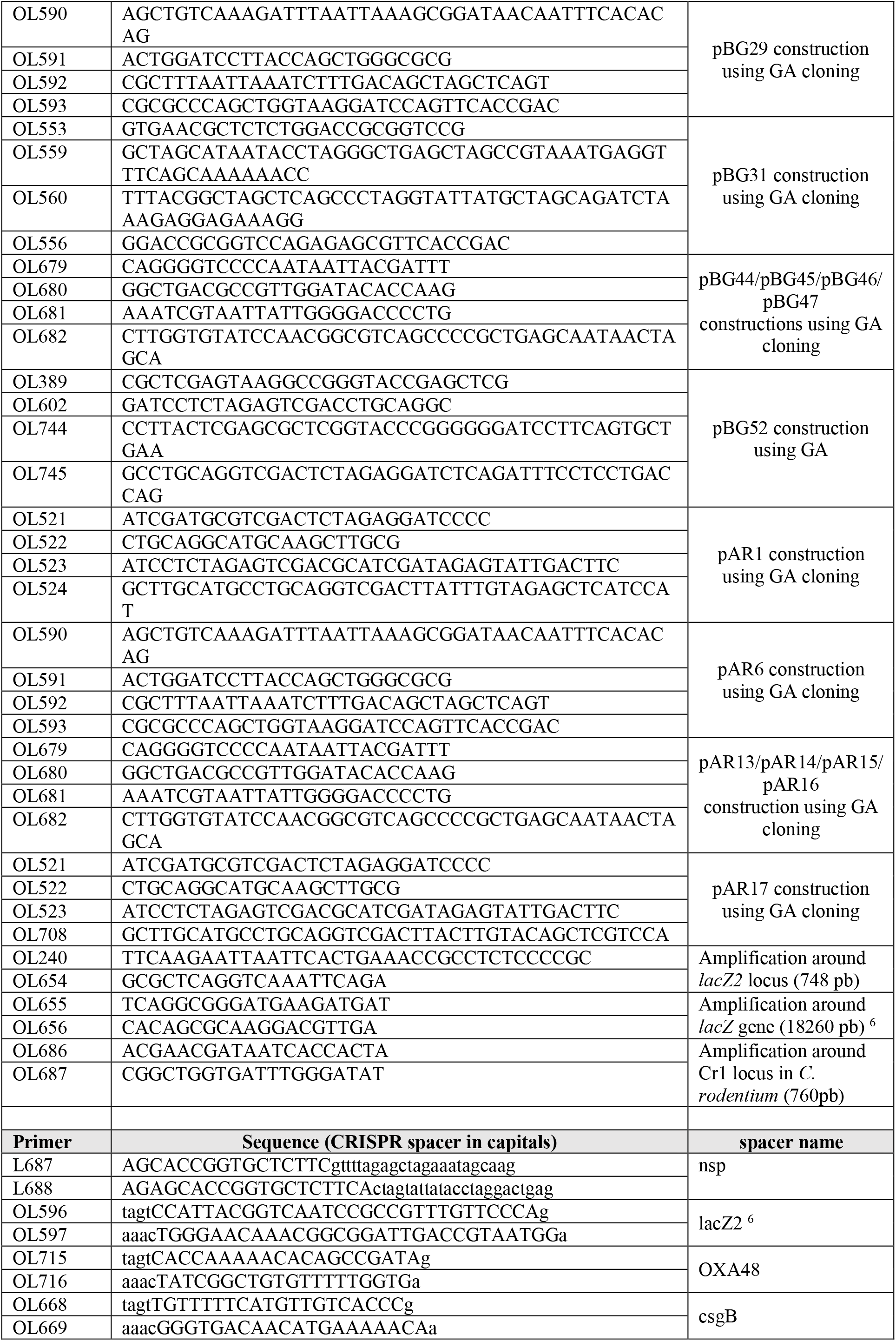
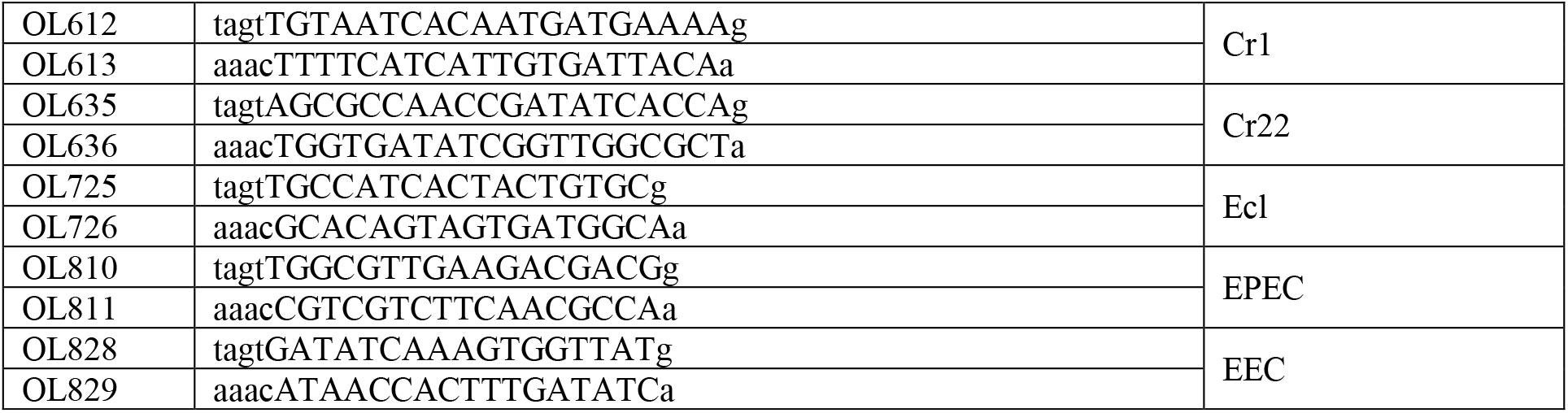
primers.

